# Degradation of Fatty Acid Export Protein1 by Rhomboid-Like Protease11 Contributes to Cold Tolerance in Arabidopsis

**DOI:** 10.1101/2023.10.26.564161

**Authors:** Annalisa John, Moritz Krämer, Martin Lehmann, Hans-Henning Kunz, Fayezeh Aarabi, Saleh Alseekh, Alisdair Fernie, Frederik Sommer, Michael Schroda, David Zimmer, Timo Mühlhaus, Helga Peisker, Katharina Gutbrod, Peter Dörmann, Jens Neunzig, Katrin Philippar, H. Ekkehard Neuhaus

**Affiliations:** Plant Physiology, University of Kaiserslautern, Erwin-Schrödinger-Str., D-67653 Kaisers-lautern, Germany; Plant Biochemistry, Faculty of Biology, Ludwig-Maximilians-Universität Munich, 82152 Planegg-Martinsried, Germany; Max Planck Institut for Molecular Plant Physiology, Wissenschaftspark Golm, D-14476 Potsdam, Germany; Molecular Biotechnology and Systems Biology, University of Kaiserslautern, Erwin-Schrödinger-Str., D-67653 Kaiserslautern, Germany; Computational Systems Biology, University of Kaiserslautern, Erwin-Schrödinger-Str., D-67653 Kaiserslautern, Germany; Institute for Molecular Physiology and Biotechnology of Plants, IMBIO, University of Bonn, Karlrobert-Kreiten-Str. 13, D-53115 Bonn, Germany; Plant Biology, Center for Human and Molecular Biology (ZHMB), Saarland University, D-66123 Saarbrücken, Germany

**Author notes:** Address correspondence to: Ekkehard Neuhaus, University of Kaiserslautern, Plant Physiology, Gottlieb Daimler-Str. 47, D-67653 Kaiserslautern, Germany. The author responsible for distribution of materials integral to the findings presented in this article in accordance with the policy described in the Instructions for Authors is: Ekkehard Neuhaus.

## Abstract

Plants need to adapt to different stresses to optimize growth under unfavorable conditions. The abundance of the chloroplast envelope located Fatty Acid Export Protein1 (FAX1) decreases after the onset of low temperatures. However, it was unclear how FAX1 degradation occurs and whether altered FAX1 abundance contributes to cold tolerance in plants. The rapid cold-induced increase in rhomboid-like protease11 (RBL11) transcript, the physical interaction of RBL11 with FAX1, the specific FAX1 degradation after RBL11 expression, and the absence of cold-induced FAX1 degradation in *rbl11* loss-of-function mutants suggest that this enzyme is responsible for FAX1 degradation. Proteomic analyses showed that *rbl11* mutants have higher levels of FAX1 and other proteins involved in membrane lipid homeostasis, suggesting that RBL11 is a key element in the remodeling of membrane properties during cold. Consequently, in the cold, *rbl11* mutants show a shift in lipid biosynthesis towards the eukaryotic pathway, which coincides with impaired cold tolerance. To demonstrate that cold sensitivity is due to increased FAX1 levels, FAX1 overexpressors were analyzed. *rbl11* and FAX1 overexpressor mutants show superimposable phenotypic defects upon exposure to cold temperatures. Our results show that the cold-induced degradation of FAX1 by RBL11 is critical for Arabidopsis to survive cold and freezing periods.

**One sentence summary:** Degradation of the inner envelope protein Fatty Acid Export1 via Rhomboid Like Protease11 represents a critical process to achieve cold and frost tolerance in Arabidopsis

## Introduction

The vast majority of vascular plants are sessile. One of their most remarkable characteristics is their ability to cope with a wide range of environmental conditions. When light intensity, temperature, or water and nutrient availability leave certain ranges, corresponding stress stimuli trigger systemic responses such as genetic, metabolic, and, to some extent, morphological changes. These processes lead to a new metabolic homeostasis that allows the plant to successfully cope with the environmental challenge (Obata and Fernie, 2012; Koevoets et al., 2016; Choudhury et al., 2017; Pommerrenig et al., 2018; Garcia-Molina et al., 2020; Wang et al., 2020).

Changes in growth temperature are usually more rapid than changes in water or nutrient availability. Accordingly, plants must rapidly initiate appropriate acclimation programs. These efficient molecular responses are essential because temperature affects the two main processes of photosynthesis, i.e., the light-driven electron transport across the thylakoid membrane and the subsequent enzyme-catalyzed Calvin-Benson cycle, in different ways. Thus, photosynthesis is markedly responsive to temperature changes, as demonstrated in several species representing a broad spectrum of CO_2_-fixing organisms (Lin et al., 2012; Mackey et al., 2013; Walker et al., 2013; Song et al., 2014).

It is well-known that the composition of membrane lipids in plant cells exhibits a dynamic remodeling after onset of low temperatures (Moellering et al., 2010; Li et al., 2015; Barnes et al., 2016; Barrero-Sicilia et al., 2017). These structural changes comprise a higher degree of desaturation and altered abundancies of different phospho-, sulfo- or galacto- lipid species, which ensure sufficient extent of membrane fluidity under unfavorable environmental conditions (Smallwood and Bowles, 2002).

In plants, lipid biosynthesis represents a complex metabolic network in which initial metabolic steps in the plastids (e.g., chloroplasts) are connected to subsequent processes at the Endoplasmic Reticulum (ER) (Li-Beisson et al., 2010; Nakamura, 2017; Hölzl and Dörmann, 2019; Lavell and Benning, 2019). Generally, for the *de novo* synthesis of both classes of lipids, namely storage lipids (tri<u>a</u>cyl-<u>g</u>lycerols, TAG) or membrane lipids (in chloroplasts mainly glyco-, phospho glycerolipids; and extraplastidic membranes, phosphoglycerolipids, sphingolipids, and sterol lipids) it is necessary that fatty acid synthesis in plastids provides acyl-chains for subsequent steps located in both, the plastid and the ER (Rawsthorne, 2002; Li-Beisson et al., 2010). Accordingly, the subsequent lipid biosynthesis takes either place via the plastid located “prokaryotic pathway” or via the ER located “eukaryotic pathway” (Roughan and Slack, 1982).

While TAG is synthesized in the ER, the biosynthesis of membrane lipid occurs in both organelles the ER and in plastids (Li-Beisson et al., 2013). During membrane lipid synthesis, the usage of fatty acids in chloroplasts or alternatively in the ER leads to different types of structural lipids. Generally, plastids represent the major site for phosphatidyl-glycerol (PG) synthesis and the exclusive site for mono- and digalactosyl- diacylglycerol (MGDG and DGDG) biosynthesis, as well as for sulfoquinovosyl- diacylglycerol (SQDG) assembly. The ER and plastids are responsible for the provision of diacyl-glycerol (DAG) backbones which serve as a precursor for all ER-borne phospholipids, including phosphatidylcholine (PC) and phosphatidylethanolamine (PE) (Hagio et al., 2002; Andersson and Dörmann, 2009; Li-Beisson et al., 2010; Lavell and Benning, 2019). (Hagio et al., 2002; Andersson and Dörmann, 2009; Li-Beisson et al., 2010; Lavell and Benning, 2019). DAG synthesis in chloroplasts is particularly pronounced under cold conditions because the Sensitive to Freezing2 (SFR2) enzyme transfers galactosyl groups from MGDG to other galactolipids, resulting in the formation of oligogalactolipids (Moellering et al., 2010; Barnes et al., 2016). Besides the plastid-produced DAG, the ER-derived DAG moieties also act - after import into plastids - as precursors for plastid lipid biosynthesis (Li-Beisson et al., 2010).

Each membrane type is defined by a characteristic lipid composition. This composition is dynamic in response to changing environmental conditions with individual lipid mixtures giving rise to specific membrane properties (van Meer et al., 2008; Li-Beisson et al., 2010; Moellering et al., 2010). For example, at low temperatures, lipid remodeling maintains membrane fluidity to prevent ion leakage, to keep carrier and receptor proteins functional or to integrate novel protective proteins (Steponkus et al., 1977; Barrero-Sicilia et al., 2017). Accordingly, plant mutants exhibiting altered activities of (i) selected fatty acid biosynthesis enzymes, of (ii) fatty acid desaturases, of (iii) lipid transfer proteins or of (iv) lipases might exhibit modified cold tolerance and photosynthesis properties (Miquel et al., 1993; Welti et al., 2002; Khodakovskaya et al., 2006; Guo et al., 2013; Gao et al., 2020; Schwenkert et al., 2023). The latter studies emphasize the impact of a proper membrane lipid remodeling after the onset of low temperatures. In fact, it has been shown for various species that low environmental temperatures lead to an upregulation of the chloroplast located lipid biosynthesis pathway (Li et al., 2015). Although several proteins involved in this process have been identified, the precise regulation responsible for this metabolic shift is still unknown.

In contrast to chloroplasts, which are the site of fatty acid *de novo* synthesis in the cell (Li-Beisson et al., 2010), the lipid biosynthesis pathway localized in the ER depends on import of fatty acids from plastids. The molecular nature of transport proteins mediating export of newly synthesized fatty acids from plastids had remained unknown for a long time (Wang and Benning, 2012). However, with the identification of the protein Fatty Acid Export Protein1 (FAX1) a first candidate was identified (Li et al., 2015) and the ability of FAX1 to promote shuttling of fatty acids across membranes has been shown in recombinant baker’s yeast cells (Li et al., 2015). Furthermore, the absence of FAX1, which resides in the plastid inner envelope membrane, leads apart from male sterility (Li et al., 2015; Zhu et al., 2020) to decreased levels of ER-derived eukaryotic lipids, while the relative content of PG, synthesized via the prokaryotic pathway, was increased. In contrast, FAX1 overexpressor lines exhibit an increased level of lipids assembled via the eukaryotic pathway, e.g., more TAG (Li et al., 2015).

The chloroplast serves as a cellular hub coordinating genetic and molecular responses required to acclimate to altered environmental conditions (Schwenkert et al., 2022). Thus, all signals and each metabolite emitted from- or received by the chloroplast must pass the inner-envelope membrane. Therefore, the inner-envelope proteome undergoes profound changes in response to the onset of stress conditions (Nishimura et al., 2016; Pottosin and Shabala, 2016; Wagner et al., 2016). Strikingly, FAX1 belongs to a group of inner envelope proteins which exhibit a proteolytic degradation after exposure to low temperatures, while other inner envelope proteins increase in their abundancy (Trentmann et al., 2020). The mechanism responsible for the decrease of FAX1 in response to cold temperatures is elusive. In addition, it is unknown whether this phenomenon represents an important molecular process required for a maximal tolerance against low temperatures and/or frost.

Chloroplasts contain more than 2000 different soluble- or membrane bound proteins (Abdallah et al., 2000) and more than 20 different proteases ensure proteome homeostasis within the organelle (Nishimura et al., 2016). So far, two major classes of proteases have been identified to cleave intrinsic inner-envelope proteins, the metallo-dependent AAA type FtsH proteases (with the isoforms FtsH 7, 9, 11 and 12) and the rhomboid-like (RBL) proteases (with the isoforms RBL 10 and 11) (Knopf et al., 2012; Wagner et al., 2016; Adam et al., 2019).

Thus, the function of FAX1 and the pronounced impact of lipid remodeling on plant acclimation to cold raise two timely questions: First, what is the mechanism responsible for the rapid decline of FAX1 upon onset of cold? Second, is the cold-induced downregulation of FAX1 a pleotropic response or is it important for low temperature tolerance? To answer both questions, we searched for an interaction of a selective protease with FAX1. It turned out, that FAX1 and RBL11 can physically interact, and that this protease is responsible for the cold-induced downregulation of FAX1. In addition, we have shown that under cold conditions, *rbl11* mutants exhibit physiological and molecular phenotypes similar to FAX1 overexpressors. Furthermore, we show that a lack of FAX1 degradation in response to cold impairs the ability of Arabidopsis plants to cope with low temperature and freezing. These findings contribute to our understanding of how plants tolerate adverse environmental conditions, one of the most impressive traits required to ensure plant productivity.

## Results

### Transcript abundance of the plastid protease *RBL11* responds to cold treatments

It was previously shown that the protein Fatty Acid Export Protein1 belongs to a small group of inner envelope proteins which are downregulated in their abundance upon onset of cold temperatures (Trentmann et al., 2020). While FtsH12 is critical for chloroplast development (Mielke et al., 2020), FtsH11 and RBL10 activity affect either, the temperature response or the membrane lipid composition, respectively (Chen et al., 2006; Wagner et al., 2011; Lavell et al., 2019). Thus, in addition to FtsH11 and RBL10, we initially focused on RBL11 (the besides RBL10 only other rhomboid-like protease at the inner envelope) which was shown to be active in fully developed mesophyll chloroplasts (Knopf et al., 2012).

To gain first insight into possible molecular interactions between the proteases and FAX1 we analyzed the transcription level of corresponding genes after transfer to 4°C (Figure 1A), the standard temperature in our laboratory to study cold effects (Patzke et al., 2019; Cvetkovic et al., 2021). Initially, transcript levels of all three envelope proteases accumulated in response to the shift towards cold growth conditions. After one week at 4°C the most pronounced *RBL11* mRNA change was detected, which had about 14-fold higher abundance when compared to the beginning of this treatment (Figure 1A). Both, *RBL10* and *FTSH11* mRNA amounts peaked at 4- or 7 times higher levels respectively, after 7 days of cold treatment versus the start of the cold period (Figure 1A). The highest levels of all three mRNAs were present after 14 days in cold conditions and from then on, these three transcript levels declined in their abundance (Figure 1A).

**Figure 1:**
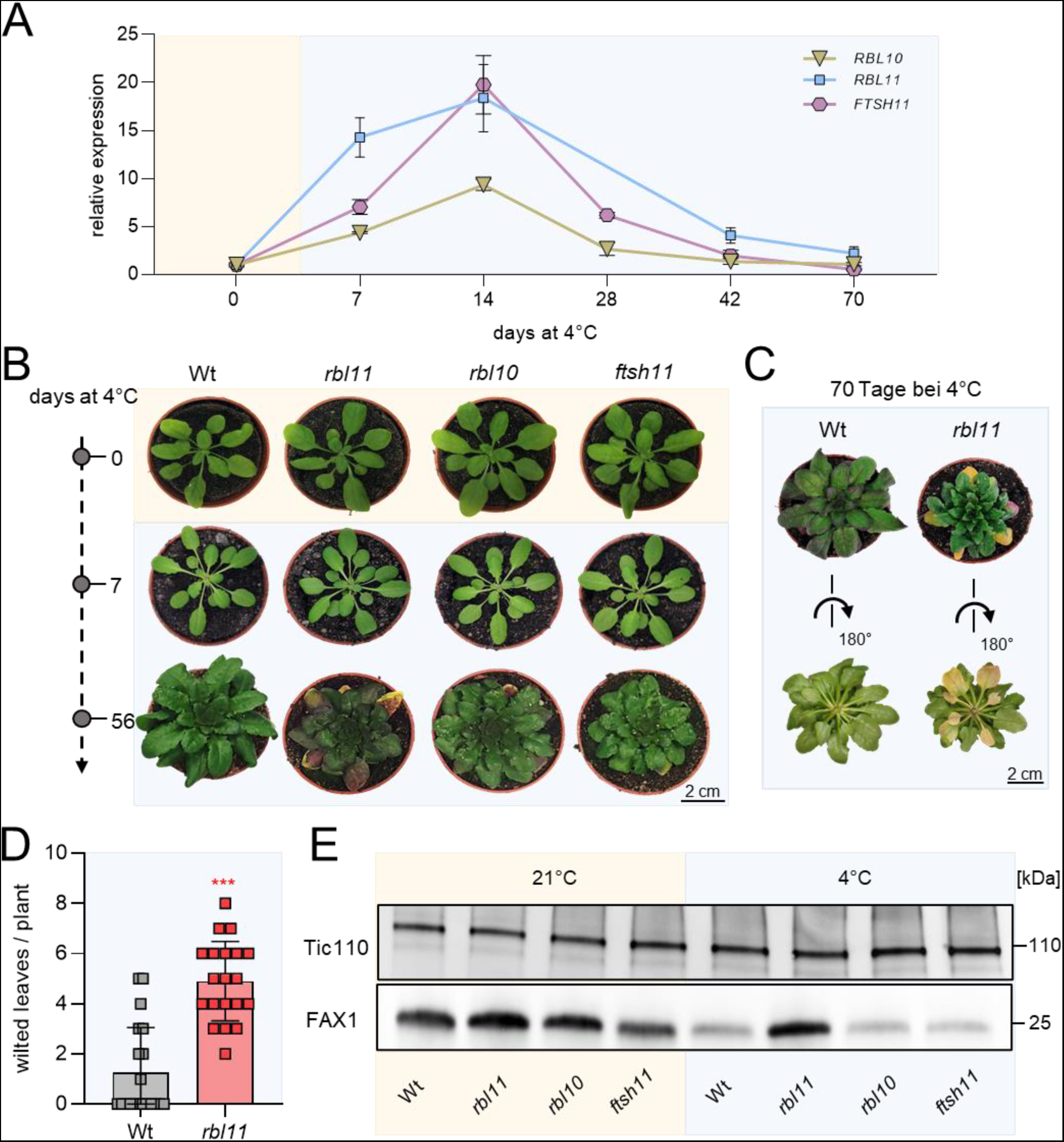
Gene expression, phenotypic and immunoblot analysis of Arabidopsis protease loss-of-function mutants *rbl11*, *rbl10* and *ftsh11* grown under standard and cold (4°C) conditions. Plants were grown under standard conditions (21°C day and night temperature, 10h day length and 120µE light intensity) for 3 weeks and then treated with cold (4°C day and night temperature, 10h day length and 120µE light intensity). A) Gene expression levels of *RBL11*, *FTSH11* and *RBL10* in wild type by qRT-PCR under standard growth conditions (0 days at 4°C) and several days during cold treatment (7; 14; 28; 42, and 70 days at 4°C). Data represent relative mean expression levels of 3 biological replicates and are normalised to standard conditions (0 days at 4°C) using UBQ as an internal control. B) Phenotypic analysis of *rbl11*, *rbl10* and *ftsh11* Arabidopsis plants. Images were taken from plants grown under standard conditions and after chilling for 7 and 56 days. C) Wt and *rbl11* rosettes were cut after 70 days of cold treatment and rotated to highlight the wilted leaves. D) Number of wilted leaves per plant in Wt and *rbl11* after 70 days of cold treatment. E) Immunoblot analysis via FAX1 antibody in isolated chloroplast envelopes of Wt, *rbl11*, *rbl10* and *ftsh11* from plants grown under standard conditions (21°C) and plants grown for 7 days at 4°C. Immunoblot detection via Tic110 antibody is used as a control and shows an equal loading of the protein samples at 3 µg per lane. Error bars in A) represent ± SEM. Error bars in D) are ± SD. Asterisks indicate significant differences between Wt and mutant using a t-test: p-value ≤0.001: *** (Supplemental file 1).

### *rbl11* mutants exhibit impaired growth at cold temperature since RBL11 controls FAX1 abundance in the cold

To search for specific responses in mutants lacking one of the three proteases, we grew wild-type and respective loss-of-function lines under either, control conditions (21°C), or for 7 or 56 days at 4°C.

When grown at 21°C, none of the mutants exhibited a phenotypic pattern distinct from wild-type plants (Figure 1B). When grown at 4°C for 7 days all genotypes showed slightly decreased chlorophyll levels, as indicated by brighter leaves (Figure 1B). However, after 56 days at 4°C all mutant plants exhibited impaired growth, i.e., smaller rosette sizes when compared to the wild type (Figure 1B). We previously showed that plants with impaired tolerance against cold temperatures exhibited increased numbers of wilted leaves when exposed to 4°C (Trentmann et al., 2020). Interestingly, after 56 days at 4°C, *rbl11* mutants exhibited wilted leaves (Figure 1B). This effect was further exacerbated in *rbl11* plants after ten weeks at 4°C (Figure 1C, pictures from top and bottom of the rosette). A quantification of this observation showed that after ten weeks at 4°C, wild types exhibited 1.25 wilted leaves/plant, while *rbl11* mutants displayed in five wilted leaves per plant on average (Figure 1D).

The observation that from all three proteases tested, the *RBL11* mRNA is the fastest responding transcript within 7 days after transfer to 4°C (Figure 1A), rendered the corresponding enzyme a prime candidate for cold-induced FAX1 degradation. To test the effect of RBL11 on FAX1 levels, we isolated chloroplast envelopes from wild types and *rbl11* loss-of-function mutant plants and conducted immunoblots using a previously established FAX1 antibody (Li et al., 2015). The parallel immunoblotting of the inner envelope protein TIC110 (Balsera et al., 2009) confirmed similar levels of total envelope proteins in each lane (Figure 1E). Interestingly, already in *rbl11* mutants grown at control temperature the FAX1 protein appeared increased by about 50% when compared to wild type plants (Figure 1E). After 7 days in cold conditions, the FAX1 protein in wild type plants decreased markedly (Figure 1E), which confirms our previous observations (Trentmann et al., 2020). Such cold-induced FAX1 degradation was almost absent in *rbl11* mutants compared to wild-type controls, while cold-induced FAX1 degradation occurred similarly in *rbl10* and *ftsh11* mutants (Figure 1E).

### *rbl11* mutants exhibit impaired frost tolerance

The observation that *rbl11* plants show an increased number of wilted leaves when grown at cold temperatures (Figure 1B-D) led us to test the freezing tolerance of this mutant. The ability of vascular plants to withstand freezing temperatures depends on a pretreatment of several days at temperatures above 0°C, called cold acclimation (Alberdi and Corcuera, 1991). To investigate whether the altered cold response in mutants affects freezing tolerance, we compared the ability of wild-types and *rbl11* plants to recover from exposure to freezing temperatures. To this end, plants were grown at 21°C and shifted to 4°C for four days to acclimate the cold. The temperature was then gradually reduced (2°C/h) until −10°C was reached. This temperature was maintained for 15h and then gradually increased (2°C/h) to 21°C before re-lighting (Trentmann et al., 2020). We assessed the phenotypic appearance of frost-treated plants after one and two additional weeks of growth at 21°C and quantified the number of wilted leaves as well as the survival rate after two weeks recovery phase.

As expected, unacclimated wild types and *rbl11* plants are unable to survive a freezing treatment (Figure 2A). As seen previously (Trentmann et al., 2020), wild-type plants are able to recover from a period of freezing with an efficiency of about 80% (Figure 2A,B) and exhibited 42% wilted leaves (Figure 2C). In contrast, *rbl11* mutants showed a significantly reduced ability to recover from freezing treatment, as only 28% of all plants survived the freezing period and more than 60% of the leaves were wilted (Figure 2A-C).

**Figure 2:**
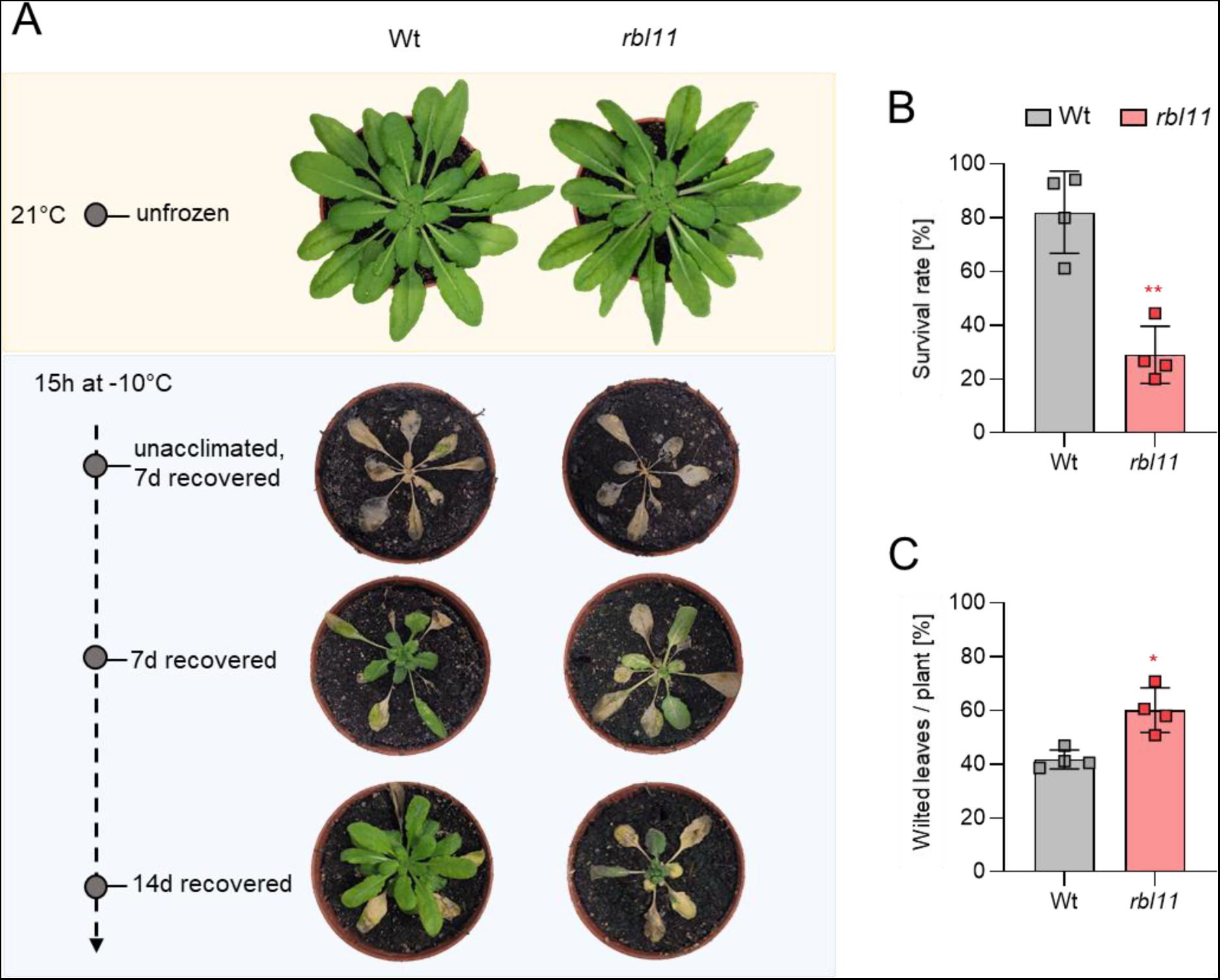
Recovery from freezing is impaired in *rbl11* loss-of-function mutants. Plants were grown under standard growth conditions for 3 weeks. Prior to freezing, the temperature was lowered to 4°C for 4 days (day and night temperature) for cold acclimation. The lowering of the temperature for freezing was done stepwise (2°C/h) and in complete darkness. The temperature for freezing was maintained at −10°C for 15 h before being gradually increased to 21°C (2°C/h). A) Representative Wt and *rbl11* plants recovered from −10°C freezing. Images were taken 7 and 14 days after freezing, from unacclimated and unfrozen (control) plants. B) Comparison of survival between Wt and *rbl11* mutants 7 days after −10°C treatment. Data represent the mean of four independent experiments with 10 to 20 plants per line and experiment. C) Quantification of wilted leaves from Wt and *rbl11* plants recovered from −10°C freezing for 7 days under standard growth conditions. Data are the mean of 4 independent experiments. Statistical differences between wild-type and overexpressor lines in B) and C) were analysed by Student’s t-test followed by Welch’s correction: p-value ≤0.05: *; p-value ≤0.01: ** (Supplemental file 1).

### RBL11 and FAX1 interact physically at the inner envelope membrane of chloroplasts

Having established that RBL11 activity is mandatory for cold-induced FAX1 degradation (Figure 1E), we were next interested to probe for a direct physical interaction between these two proteins. To this end, we exploited the Bimolecular Fluorescence Complementation (BiFC) analysis (Kerppola, 2008) based on the association of complementary yellow fluorescent protein (YFP) fragments fused to putative partner proteins. Only when both partners are in close proximity, a functional fluorescent protein can be formed, indicating the protein-protein interaction and its subcellular location within the living cell.

We infiltrated *Nicotiana benthamiana* leaves with the constructs FAX1:YFP^C-n^and RBL11:YFP^C-c^ for synthesis of the respective YFP fragment proteins. Co-expression of both constructs resulted in yellow fluorescing spots in epidermis cells (Figure 3A), indicating that both proteins interact in corresponding plastids. To further visualize the complex formation of both proteins at the chloroplast envelope, we infiltrated *N. benthamiana* leaves with both constructs and subsequently isolated mesophyll protoplasts. The combined expression of FAX1:YFP^C-n^ and RBL11:YFP^C-c^ led to a yellow fluorescence of chloroplasts (Figure 3B). The ring-shaped fluorescence (Figure 3B) resembles the YFP fluorescence emitted by other inner envelope associated YFP fusion proteins (Witz et al., 2014; Patzke et al., 2019). To validate the BiFC data we performed further control experiments using inner envelope associated protease FtsH11. In contrast to the co-expression of FAX1:YFP^C-n^ and RBL11:YFP^C-c^, the infiltration of the plasmids with FAX1:YFP^C-n^ and FTSH11:YFP^C-c^ did not yield fluorescence complementation (Figure 3C,D). In addition, the combined infiltration of the constructs FTSH11:YFP^C-n^ and RBL11:YFP^C-c^ also did not give rise to a fluorescence signal (Figure 3E,F). These two independent control experiments indicate that protein-protein interaction at the inner envelope does not occur by chance, highlighting the specificity of the RBL11 and FAX1 interaction (Figure 3A,B).

**Figure 3:**
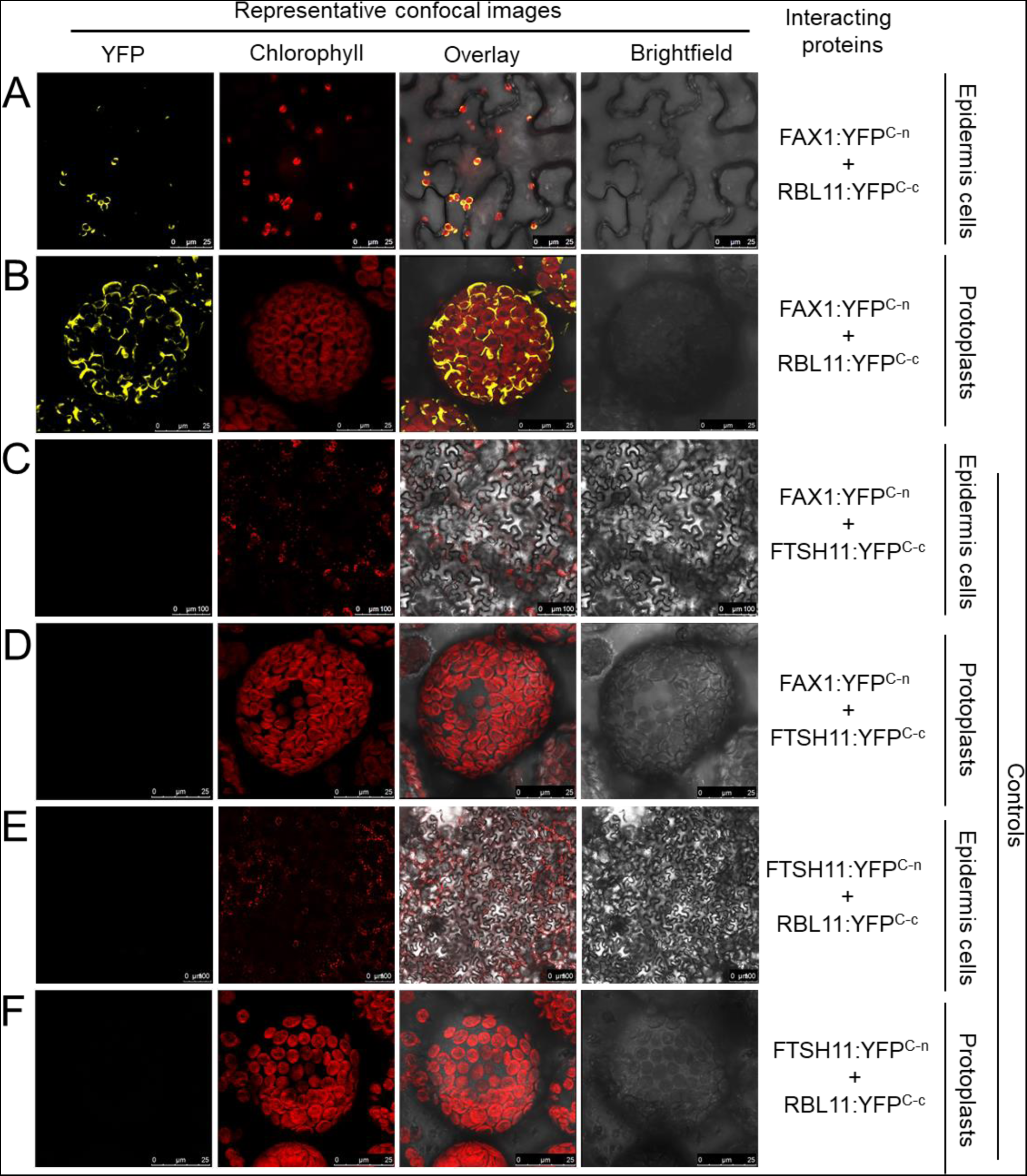
Probing of putative RBL11 targets. Constructs of full-length coding sequences were fused upstream to the C-terminus of yellow fluorescent protein (YFP) and transiently expressed in *Nicotiana benthamiana* leaves by agroinfiltration. YFP and chlorophyll fluorescence signals were recorded 4 days after infiltration in epidermis cells as well as in isolated protoplasts by confocal microscopy.

### Induction of *RBL11* leads to a specific degradation of FAX1

In contrast to wild types, *rbl11* plants are almost unable to degrade FAX1 after the onset of cold (Figure 1E). To further support our hypothesis that RBL11 is responsible for cold-induced FAX1 degradation, we generated a stable RBL11-HA (HA = hemagglutinin) over-expressor line in which the recombinant *rbl11* gene was placed under the control of a dexamethasone (DEX)-inducible promoter (Aoyama and Chua, 1997) (Figure 4A). In the presence of DEX, the RBL11-HA protein is synthesized in mutant leaf discs at 21°C and at 4°C (Figure 4B,C), and neither DEX-induced overexpression nor the C-terminal HA tag alters the membrane localization of RBL11 (Figure 4B,C). Subsequent enrichment of total leaf membranes, followed by immunoblot analysis using the FAX1-specific antibody, showed that RBL11 overexpression does not lead to FAX1 degradation from leaf discs incubated with DEX at 22°C (Figure 4D), which might be due to some post-translational modification required for FAX1 activity. In contrast, DEX incubation at 4°C leads to FAX1 degradation (Figure 4D, note that extraction of total membrane proteins from whole membranes enriched from cold treated leaf discs generally resulted in a higher abundance of FAX1 (Figure 4D, left and right panels).

**Figure 4:**
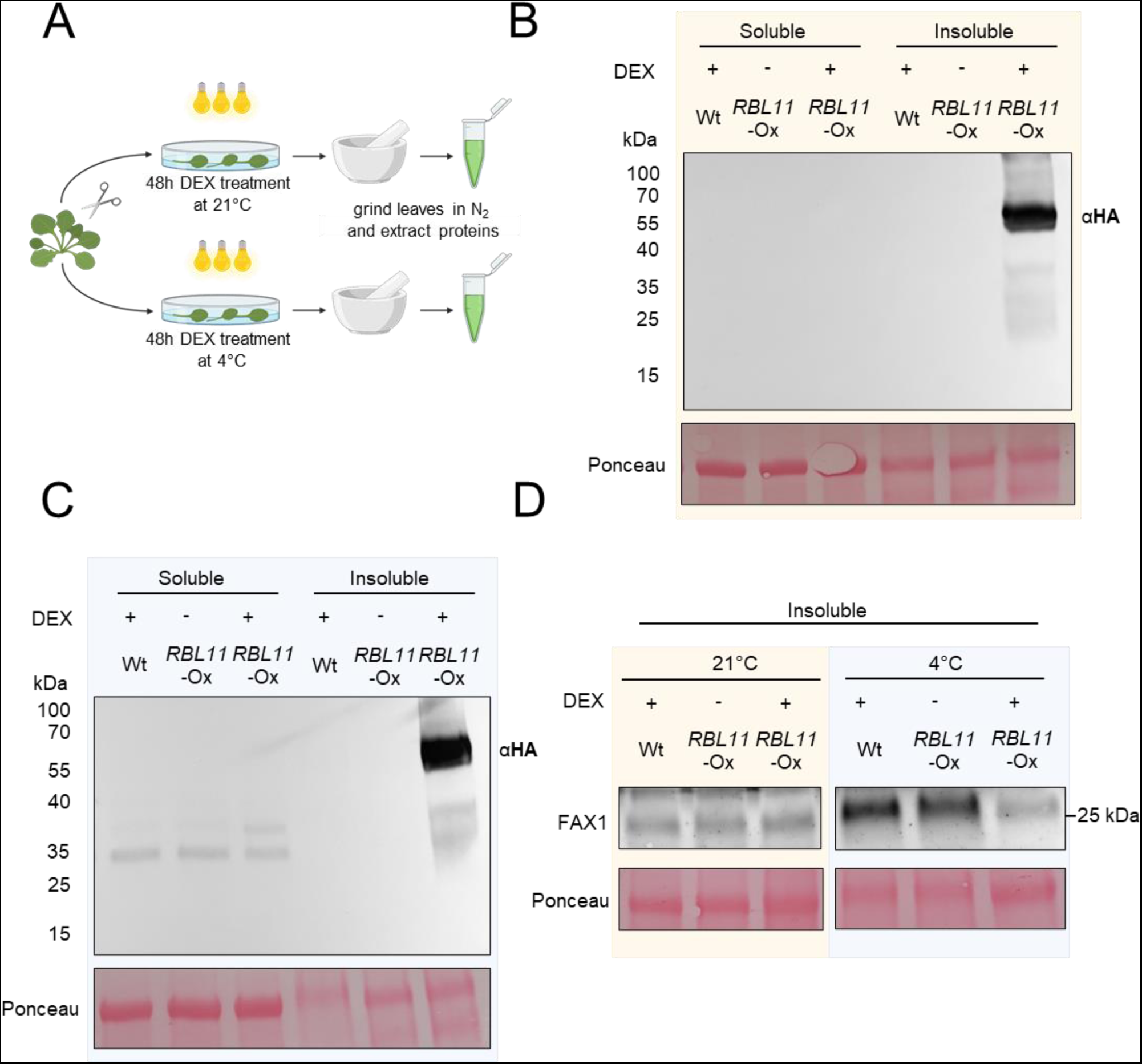
Cold induced degradation of FAX1 by RBL11 in Arabidopsis. A) Scheme of experimental setup. B) Immunoblot analysis via HA antibody of soluble and insoluble (membrane) protein extracts fom wildtype and transgenic plants expressing RBL11 tagged with a HA epitope (∼ 32 kDa) under the control of an 35S promotor after induction with dexamethasone for 48h at 21°C. C) Immunoblot detection via HA antibody of soluble and insoluble (membrane) protein extracts fom wildtype and transgenic plants expressing RBL11 tagged with a HA epitope (∼ 67 kDa) under the control of an 35S promotor after induction with dexamethasone for 48h at 4°C. Note that the size of the protein (∼ 67 kDa) comes from the presence of an additional biotinylase (∼35 kDa), which was not relevant in this experiment. D) Immunoblot of insoluble protein extracts from Wt and and transgenic plants overexpressing *RBL11* under the control of an 35S promotor after induction with DEX for 48h at 21°C or 4°C with a FAX1 antibody. Please note that the extraction of whole membranes enriched from cold treated leaves generally resulted in a higher abundance of FAX1 and is not compareable with results from leaves treated at 21°C. Ponceau staining in B), C) and D) is representing equal loading of protein samples with 18 µg per lane.

### The membrane lipid composition of *rbl11* mutants indicates a shift to the eukaryotic biosynthesis pathway under low temperature

The abundance of FAX1 strongly decreases after transfer of wild type to cold conditions (Figure 1E and Trentmann et al. 2020). Because the absence of RBL11 prevents cold-induced FAX1 degradation (Figure 1E) we aimed to determine changes in lipid levels under these growth conditions. For this purpose, wild-type and *rbl11* mutant plants were first grown for 4 weeks under control conditions and then either, for further 14 days at 21°C or at 4°C, prior to extraction and quantification of leaf lipids.

Growth of plants at 21°C led to lower leaf levels of the major glycolipids monogalactosyl-diacylglycerol (MGDG) and digalactosyl-diacylglycerol (DGDG) in *rbl11* plants compared to wild types (Figure 5). In contrast, the levels of sulfoquinovosyl-diacylglycerol (SQDG) and the phospholipids phosphatidic acid (PA), phosphatidylserine (PS), phosphatidylinositol (PI), phosphatidylglycerol (PG), phosphatidylethanolamine (PE) and phosphatidylcholine (PC) were similar in leaves from the two plant lines (Figure 5). These absolute amounts summed up to 230 nmol mg^-1^ DW of phospho- and glycolipids in *rbl11* mutants and 274 nmol mg^-1^ DW of phospho- and glycolipids in wild-type plants (Figure 5A, inset). However, after recalculating the data from total levels into mol% no obvious changes in the relative individual lipid species between *rbl11* mutant and wild type plants were found (Figure 5B). Interestingly, growth for 14 days under cold temperature conditions led to higher total lipid levels in the *rbl11* plants (Figure 5C inset). This increase of total lipids in the *rbl11* mutants is due to higher levels of MGDG, DGDG and SQDG, but also to higher levels of all phospholipids (Figure 5C). Furthermore, after recalculating the data into mol% it appeared that growth at 4°C did not only lead to higher total lipid contents in *rbl11* plants, but that especially the relative proportions (in mol%) of the phospholipid species PA, PI, PE and PC to the total lipids was increased in the *rbl11* plants, while proportions of MGDG and DGDG are decreased (Figure 5D).

**Figure 5.**
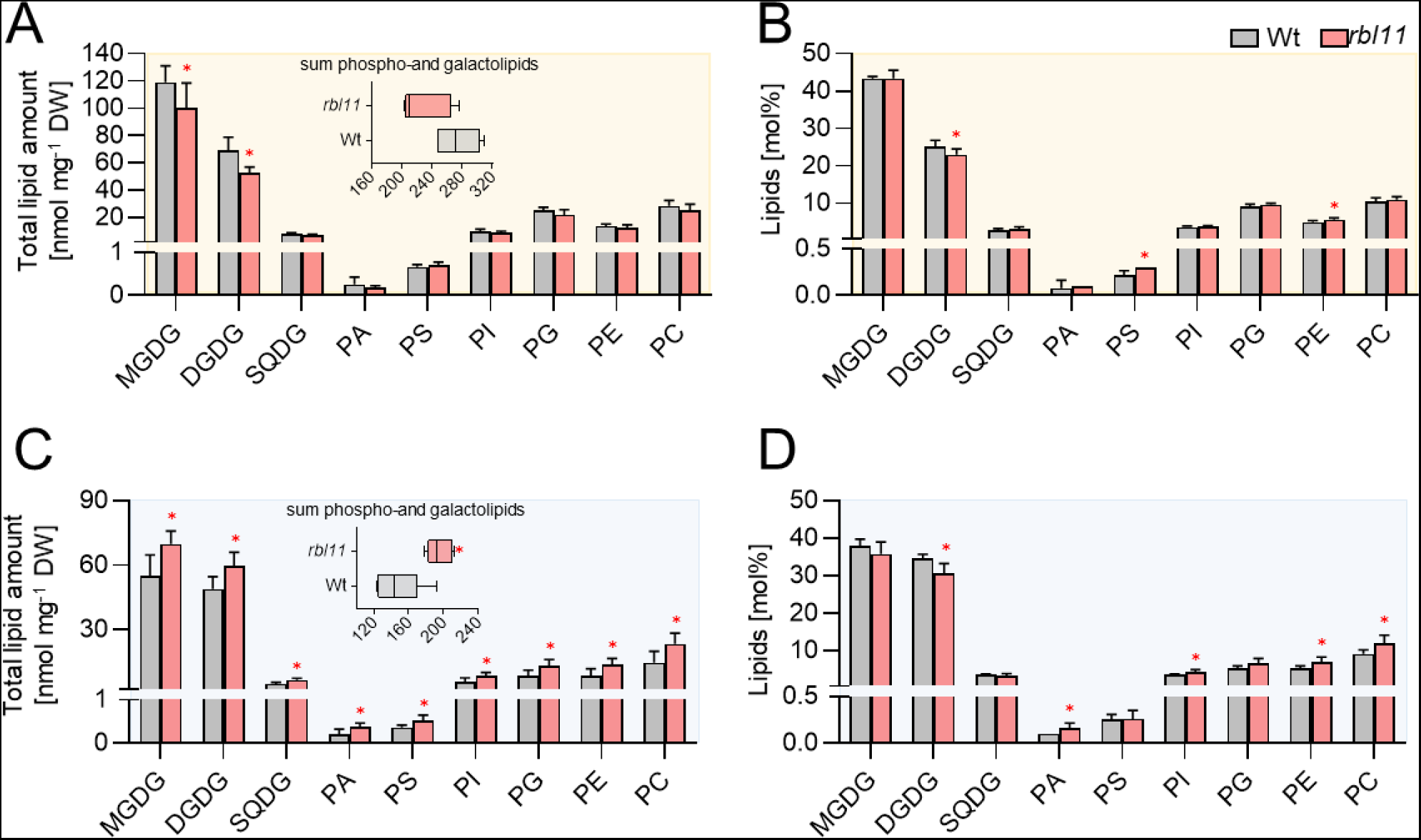
Analysis of galacto- and phospholipids in Wt and *rbl11* loss-of-function mutants. Changes in A) total contents and B) relative amounts of different galacto- and phospholipids in rosette leaves of 3-week-old Arabidopsis plants grown under standard conditions. The box plot in A) shows changes in the sum of the measured total lipid contents. Differences in C) total lipid contents and D) relative galacto- and phospholipid amounts in plants grown under standard conditions for 3 weeks before lowering the temperature to 4°C for 14 days. The box plot in C) shows the changes in the sum of the measured total lipid contents. Data represent the mean of 5 plants per row. Error bars indicate ± SD. Significance of differences between wild-type and mutant was analysed by Student’s t-test: p-value ≤0.05:* (Supplemental file 1).

### Identification of other putative RBL11 protease substrates

By analyzing the envelope proteome of *rbl10rbl11* double mutants an overrepresentation of non-degraded envelope proteins compared to wild type emerged (Knopf et al., 2012). However, this type of analysis did not allow for the identification of RBL11-specific substrates. Thus, we compared the envelope proteome from wild type plants with that of *rbl11* single mutants (Table 1, Suppl. Table 1, Suppl. Table 2, and Suppl. Table 3). Since we were especially interested in RBL11 substrates degraded after onset of cold temperatures, these analyses were carried out on plants grown at 21°C or at 4°C (Table 1).

**Table 1.** The table shows the 15 top hits of proteins that are increased by more than 30% in the mutant at low temperature. For each protein a two-way ANOVA was performed, which means that only proteins with a significance level below 0.05 were considered. The qValue_Interaction Genotype Temperature indicates that there is a temperature effect that differs between mutant and wild type. All proteins were annotated by name, gene locus and description. The proteins written in bold and in italics are increased under standard conditions as well as after 7 days at 4°C (see Suppl. Table 1) to the described selected criteria (Abundance > 30%; qValue < 0.05). FAX1 was labeled with an asterisk because the protein is not within the top hits, but also fulfills the selected criteria.

| Protein | Gene locus | Description | Log2FC<br>*rbl11* / Wt | qValue_Interaction<br>Genotype Temperature |
| --- | --- | --- | --- | --- |
| **TGD5** | At1g27695 | Encodes a small glycine rich protein that is localized to the chloroplast envelope and is a component of the ER to plastid lipid trafficking pathway. | 4.54 | 0.033 |
| **PORA** | At5g54190 | light-dependent NADPH:protochlorophyllide oxidoreductase A | 4.08 | 0.030 |
| HCF107 | At3g17040 | Involved in regulating plastidial gene expression and biogenesis | 1.56 | 0.002 |
| **unknown** | At3g28220 | TRAF-like family protein, contains MATH domain | 2.73 | 0.044 |
| KIN14A | At5g10470 | Kinesin-like protein required for chloroplast movements and anchor to the plasma membrane | 2.18 | 0.032 |
| **DVR** | At5g18660 | Encodes a protein with 3,8-divinyl protochlorophyllide a 8-vinyl reductase activity | 1.85 | 0.031 |
| **unknown** | At1g08530 | Chitinase-like protein | 1.35 | 0.019 |
| unknown | At5g19850 | Alpha/beta-Hydrolases superfamily protein | 1.47 | 0.027 |
| **unknown** | At1g64850 | Calcium-binding EF hand family protein | 1.22 | 0.015 |
| unknown | At5g39410 | Saccharopine dehydrogenase | 1.06 | 0.018 |
| unknown | At2g43630 | Nucleus envelope protein | 0.48 | 0.004 |
| TGD2 | At3g20320 | Encodes a permease-like component of an ABC transporter involved in lipid transfer from ER to chloroplast. A phosphatidic acid-binding protein. | 0.98 | 0.022 |
| DEGP2 | At2g47940 | Encodes DegP2 protease | 0.83 | 0.015 |
| SPPA | At1g73990 | Encodes a light-inducible chloroplast protease complex | 0.78 | 0.015 |
| FAX1* | At3g57280 | involved in fatty acid and lipid homeostasis and likely functions as a fatty acid transporter that exports fatty acids from the plastid | 0.38 | 0.028 |

Putative substrates of RBL11 are expected to be more abundant in *rbl11* mutants when compared to wild-type plants. In a previous study on proteins accumulating in the inner envelope of *rbl10rbl11* double mutants, the TRAF-like family protein containing a MATH domain was also found to accumulate in the double mutant (Knopf et al., 2012). Similarly, the TRAF-like family protein accumulated in *rbl11* single mutants (Suppl. Table 1) indicating that this protease is responsible for its degradation. Interestingly, the abundance of the protein TGD5, which is involved in ER to plastid lipid transfer (Fan et al., 2015), is also higher in *rbl11* mutants than in wild types, and this effect is independent of the environmental temperature (Suppl. Table 1 and Table 1). Besides TGD5, the abundance of another TGD protein, namely TGD2, is also higher in *rbl11* mutants when exposed for 4 days to 4°C (Table 1). In addition, the chloroplast located protease systems DegP2 and SPPA are also more abundant in cold treated *rbl11* mutants when compared to wild type (Table 1). Similarly, the inner envelope associated NADPH:protochlorophyllide reductase PORA (Barthélemy et al., 2000) appears, besides the TGD components and FAX1, as a further RBL11 substrate (Suppl. Table 1), especially after onset of cold temperature (Table 1). It seems likely that the decrease of some proteins in *rbl11* chloroplasts under warm or cold conditions represents pleiotropic reactions of the mutant induced by altered FAX1 abundance (Suppl. Table 2 and Suppl. Table 3). Nevertheless, among these proteins, we also found proteins associated with lipid metabolism, e.g., LACS9, representing a major long chain acyl-CoA synthetase (Schnurr et al., 2002), or BASS1, representing a sodium/pyruvate cotransporter (Furumoto et al., 2011) (Suppl. Table 2 and Suppl. Table 3).

### *FAX1* overexpressor lines exhibit markedly increased levels of FAX1 protein

The above data indicate a molecular interrelation between increased FAX1 protein abundance (due to an absence of RBL11) and the levels of membrane lipids and Arabidopsis cold tolerance (Figures 1 and 3). This encouraged us to analyze the impact of increased FAX1 abundance on plant cold tolerance. *rbl11* mutants are not conducive to this, because the absence of this protease does not exclusively affect FAX1 levels, but also the abundance of other envelope associated proteins (Table 1). To avoid the study of pleiotropic effects, we decided to investigate the cold tolerance of Arabidopsis mutants with constitutively increased FAX1 levels (Li et al., 2015).

Our observations that *rbl11* loss-of-function mutants have increased FAX1 levels under cold conditions (Figure 1E) and that directed synthesis of RBL11 leads to a decrease in FAX1 in the cold (Figure 2) provide evidence that the RBL11 protease is responsible for this process. Nevertheless, for a comprehensive understanding of FAX1’s impact on the cold response, it is mandatory to study *FAX1* transcript levels after transfer to cold conditions. In fact, the observation that *FAX1* mRNA rises markedly in wild-type plants after exposure to 4°C (Figure 6A) supports our conclusion that RBL11 is responsible for the cold-induced decrease of the level of the FAX1 protein (Figure 1F).

**Figure 6:**
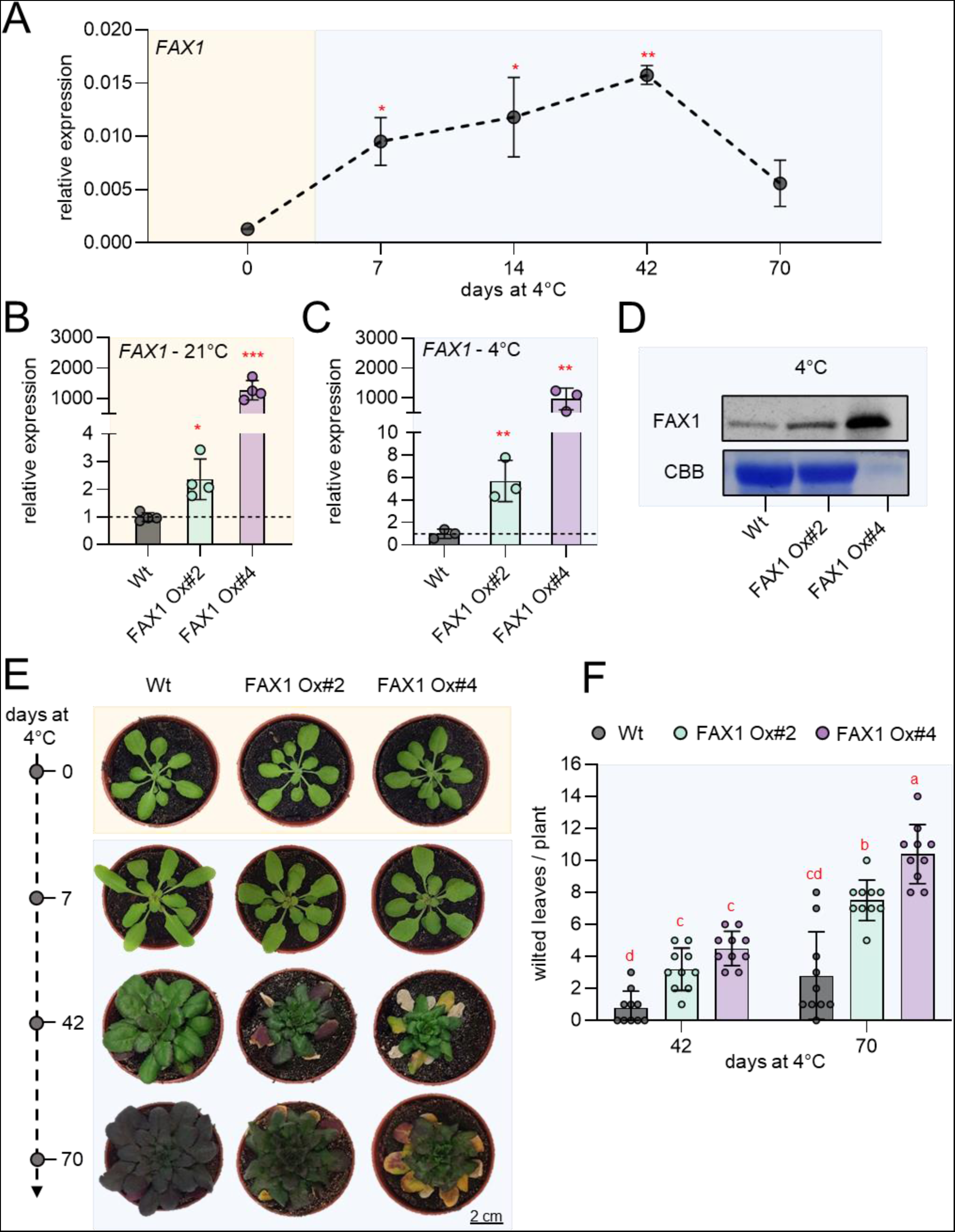
Gene expression, immunoblot and phenotypic analysis of two independent Arabidopsis fatty acid export protein 1 (FAX1) overexpression lines (FAX1 Ox#2 and FAX1 Ox#4) and wild-type (Wt) plants grown under standard and cold (4°C) conditions. Plants were grown under standard conditions (21°C day and night, 10h day length and 120µE light intensity) for 3 weeks and then treated with cold (4°C day and night, 10h day length and 120µE light intensity). A) Expression of FAX1 by qRT-PCR under standard growth conditions (0 days at 4°C) and several days during cold treatment (7; 14; 42, and 70 days at 4°C). Data represent relative mean expression levels of 3 biological replicates and are normalised to standard conditions (0 days at 4°C) using UBQ as an internal control. Relative expression of FAX1 by qRT-PCR under B) standard growth conditions and C) 7 days after cold treatment. Data represent relative mean expression levels and are normalised to the wild type using UBQ as an internal control. D) Immunoblot analysis of FAX1 in crude extract from plants grown at 4°C for 7 days. The image of the Coomassie stained gel (CBB) shows equal loading of protein samples from Wt and FAX1 Ox#2 with 30µg per lane. The amount of FAX1 Ox#4 protein is reduced to 3µg. E) Rosette phenotype of representative Wt, FAX1 Ox#2 and FAX1 Ox#4 under standard growth conditions (0 days at 4°C) and after cold treatment (7; 42, and 70 days at 4°C). F) Number of wilted leaves per plant after 42 and 70 days of cold treatment. Error bars in A) represent ± SEM. Error bars in B), C) and F) are ± SD. Statistical differences between wild-type and overexpressor lines in A), B) and C) were analysed by Student’s t-test: p-value ≤0.05: *; p-value ≤0.01: **; p-value ≤0.001: ***. Letters above error bars in F) indicate significant differences by two-way ANOVA followed by Tukey’s test (p<0.05; (Supplemental file 1)).

Previously, we generated two independent FAX1 overexpressing lines, FAX1 Ox#2 and FAX1 Ox#4, both containing the *FAX1* gene under the control of the constitutive cauliflower 35S promotor (Li et al., 2015). So far, it has only been shown that these mutant lines have both, increased *FAX1* mRNA and increased FAX1 protein levels when grown at ambient temperature (Li et al., 2015). We verified that the *FAX1* mRNA accumulation is a stable feature when FAX1 overexpressor mutants are cultivated at cold temperature. For this purpose, the *FAX1* mRNA levels were quantified in plants grown under the standard temperature of 21°C with those grown for one week at 4°C. The two FAX1 overexpressor lines exhibited a substantial accumulation of *FAX1* mRNA when compared to wild-type plants. Under warm temperature, FAX1 Ox#2 plants contained about 2.1-fold higher *FAX1* mRNA levels as present in the wild type, while FAX1 Ox#4 plants contained even 1170-fold higher *FAX1* mRNA (Figure 6B). This marked difference of *FAX1* mRNA in the two overexpressors concurs with previous observations (Li et al., 2015). After one week of growth at 4°C, FAX1 Ox#2 plants contained about 5.7-fold higher *FAX1* mRNA levels, and FAX1 Ox#4 plants contained more than 970-fold higher *FAX1* mRNA levels than present in the wild type (Figure 6C). Thus, even at 4°C, both FAX1 overexpressor lines exhibited higher *FAX1* mRNA than present in the wild type.

It was shown that both FAX1 overexpressors contain higher levels of FAX1 protein when grown at 21°C (Li et al., 2015). To confirm that also under cold conditions, FAX1 overexpressor lines exhibit higher FAX1 protein levels than present in wild types, we conducted a immunoblot analysis (Li et al., 2015). After one week of growth at 4°C, the relative FAX1 protein level in both overexpressor plants was higher than observed in wild type plants (Figure 6D), and a corresponding quantification experiment revealed that Ox#2 plants contained about 3-fold more FAX1 protein than present in wild type, while Ox#4 plants contained about 115-fold more FAX1 protein (Figure 6D, please note: loaded protein extracted from Ox#4 plants was 1:10 diluted when compared to proteins extracted from wild types and Ox#2 mutants).

### Similar to *rbl11* mutants, FAX1 overexpressor plants exhibit decreased chilling and frost tolerance

The data above indicate that *rbl11* mutants exhibit increased cold sensitivity (Figure 1B, C). Because *rbl11* mutants revealed higher FAX1 levels than present in wild types it was interesting to analyze whether the two lines constitutively expressing the *FAX1* gene also show altered response to low temperatures.

Therefore, we cultivated all plants at ambient temperature (21°C) prior to cultivation at 4°C. Subsequently, we inspected morphologic plant appearance and the number of wilted leaves in the respective growth phase. After 7 days at 4°C none of the three plant lines developed wilted leaves (Figure 6E). However, after 42 days at 4°C, and even more pronounced after 70 days at 4°C, leaves from both FAX1 overexpressors gradually wilted more rapidly than observed in the wild type (Figure 6E).

To quantify this observation we counted the wilted leaves as a proxy for chilling sensitivity (Trentmann et al., 2020). After 42 days at 4°C wild type plants exhibited on average one wilted leaf while FAX1 overexpressor Ox#2 exhibited in average three wilted leaves, and Ox#4 mutants showed in average 4.5 wilted leaves (Figure 6F). After growth for ten weeks at 4°C, wild type plants displayed on average 2.8 wilted leaves per plant while FAX1 overexpressors Ox#2 and Ox#4 exhibited on average 7.5 and 10.4 wilted leaves, respectively (Figure 6F).

As done for *rbl11* plants (Figure 2), we compared the frost tolerance of FAX1 over-expressor plants and wild types. In contrast to both FAX1 overexpressors, wild types which survived the post-freezing phase restored efficient growth within the next 14 days, as indicated by a larger rosette size and less wilted leaves when compared to over-expressor mutants (Figure 7A). FAX1 overexpressor plants able to recover from freezing exhibited about 80% of wilted leaves, while wild types only exhibited about 46% of wilted leaves (Figure 7A, B). With about 87% the survival rate of wild types reached a value similar to previous observations (Trentmann et al., 2020). In contrast, only 57% of the FAX1 Ox#2 plants survived this stress, and only 43% of FAX1 Ox#4 plants recovered from frost (Figure 7C).

**Figure 7:**
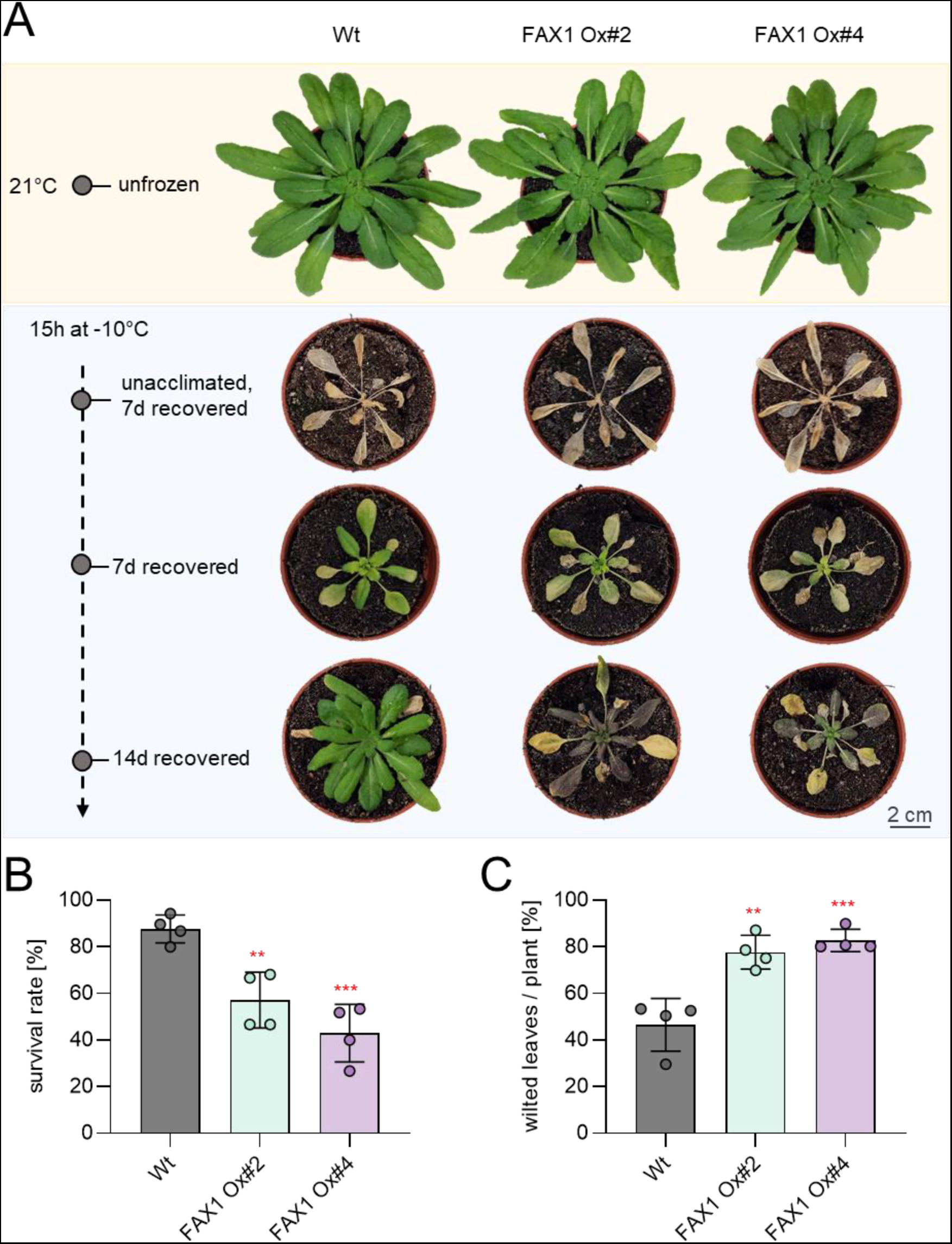
Recovery from frost is impaired in FAX1 overexpression mutants. Plants were cultivated for 3 weeks under standard growing conditions. Before freezing the temperature was lowered to 4°C for 4 days (day and night temperature) for cold acclimation. Lowering of the temperature for freezing was done stepwise (2°C/ h) and in completely dark. −10°C was kept for 15 h before a stepwise temperature raising to 21°C (2°C/h). A) representative Wt and FAX1 overexpression plants recovered from −10°C freezing. Pictures were taken 7 and 14 days after freezing, from unacclimated and unfrozen (control) plants. B) Comparison of survival rate between Wt and FAX1 overexpression mutants 7 days after −10°C treatment. Data represent the mean value from four independent experiments with 11 to 15 plants per line and experiment. C) Quantification of wilted leaves from Wt and FAX1 overexpressor plants recovered for 7 days from −10°C freezing under standard growing conditions. Data represent the mean value of 4 independent experiments. Statistical differences between wildtype and the overexpressor lines in B) and C) was analyzed by Student’s t-test: p-value ≤0.01: **; p-value ≤0.001: *** (Supplemental File 1).

### Similar to *rbl11* mutants, lipid biosynthesis in FAX1 overexpressors is shifted to the eukaryotic biosynthesis pathway under low temperature

Under cold conditions, *rbl11* mutants exhibited a shift towards ER-synthesized membrane lipids when compared to their corresponding wild type (Figure 5). To check whether this response is due to high FAX1 protein abundance, we analyzed the lipid composition in FAX1 overexpressors. For a first overview of cold effects on membrane lipid homeostasis in FAX1 overexpressor plants, we grew wild types and the representative FAX1 Ox#4 line (Li et al., 2015) under either control conditions, or for two weeks at 4°C. Subsequently, leaf lipids were extracted and quantified via mass-spectrometry.

When grown at 21°C, the total levels (nmol mg^-1^ DW) and the relative proportions (mol%) of the glycolipids MGDG, DGDG and SQDG, or of the phospholipids PA, PS, PI, PG, PE and PC were similar in leaves from wild types and FAX1 Ox#4 plants (Supplemental Figure 1A, B and insert). In contrast, after two weeks at 4°C FAX1 overexpressor plants contained higher levels of MGDG and DGDG when compared to wild types (Supplemental Figure 1C). In addition, the levels of PS-, PG-, PE- and PC FAX1 Ox#4 plants appeared to be slightly higher than in corresponding wild types (Supplemental Figure 1C). This increase led in sum to 158 nmol mg^-1^ DW of membrane lipids in FAX1 Ox#4 plants, while wild types contained only 125 nmol mg^-1^ DW membrane lipids (insert Supplemental Figure 1C). Similar to the situation at 21°C, the proportions of all lipid species in both plant lines are nearly identical (Supplemental Figure 1D).

Since FAX1 overexpressor plants revealed impaired cold tolerance after long exposure to low temperatures, and because the observed changes in lipid composition of FAX1 overexpressors after two weeks of growth at cold temperature appeared in parts to be moderate (Supplemental Figure 1), we extracted lipids from plants exposed for ten weeks to 4°C (Figure 8). Similar to plants exposed to only two weeks at 4°C (Supplemental Figure 1), both FAX1 overexpressor lines exhibited increased total levels of the galactolipids MGDG and DGDG, and also of the phospholipids PI, PG, PE and PC (Figure 8A). The increase of the two groups of lipids led to an overall higher level of total membrane lipids in FAX1 overexpressor lines (FAX1 Ox#2, 205 nmol mg^-1^ DW; FAX1 Ox#4, 188 nmol mg^-1^ DW; wild type, 125 nmol mg^-1^ DW; (insert Figure 8A). The levels of the low abundant glycolipid SQDG and the phospholipid PS were not altered in FAX1 overexpressors, when compared to wild type (Figure 8A). Although total amounts of glycolipids in both FAX1 overexpressors were higher when compared to wild types (Figure 8A), the relative contribution (mol %) of MGDG, DGDG and SQDG in FAX1 Ox#2 and #4 to total membrane lipids was similar to wild type (Figure 8B), while the relative contribution of the phospholipids PG, PE and PC in FAX1 Ox#2 and #4 to total membrane lipids was increased, when compared to the wild type (Figure 8B). The most pronounced alterations of phospholipids were noted for PC and PE (Figure 8B), which were 40% higher in FAX1 overexpressors than in wild type plants (Figure 8B).

**Figure 8:**
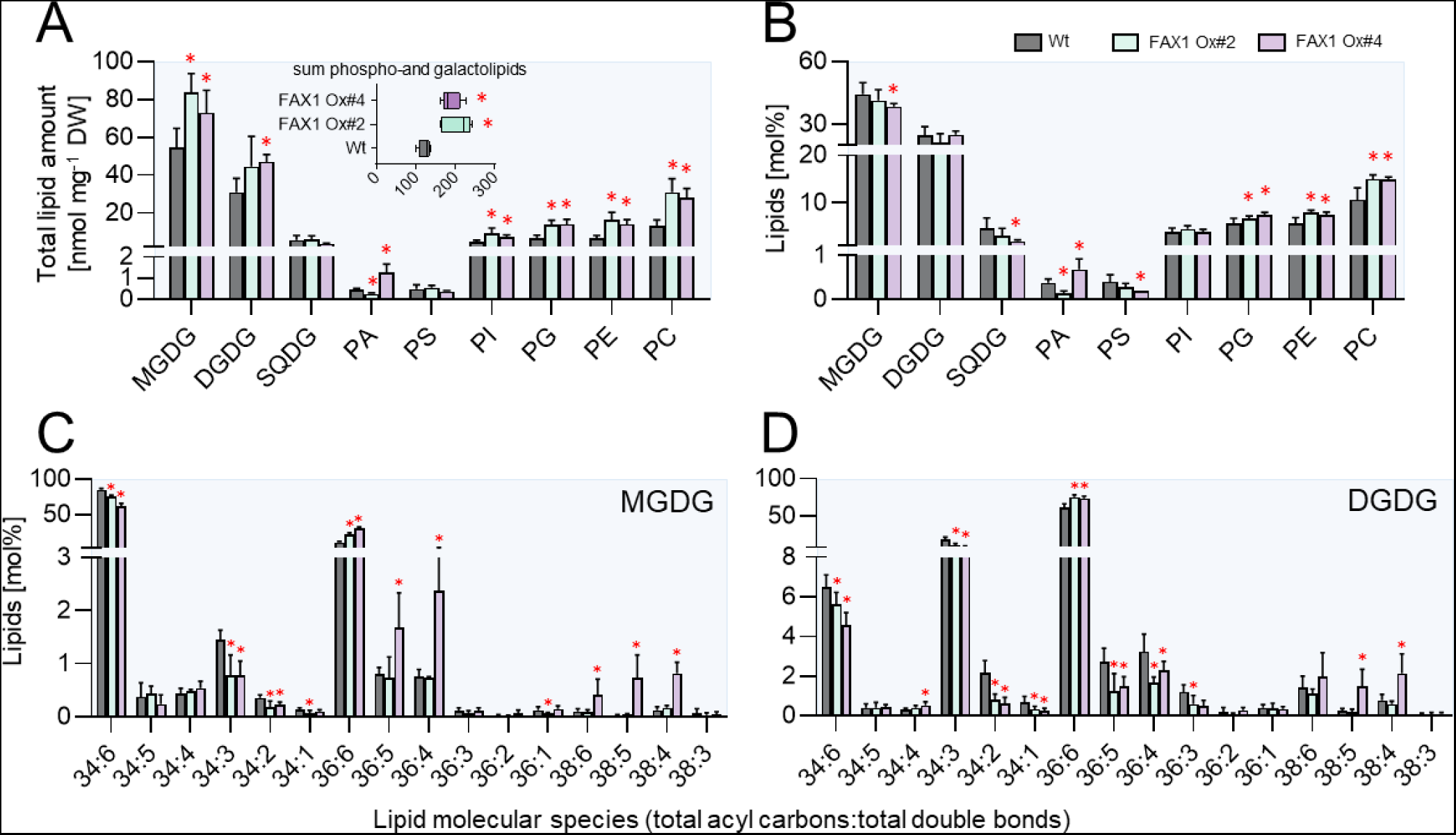
Lipid analysis of Wt and FAX1 overexpression lines. Galacto- and phospholipids as well as galactolipid molecular species were determined in leaves of plants grown for 3 weeks under standard conditions before lowering the growth temperature to 4°C for 10 weeks. Changes in A) total contents and B) relative amounts of different galacto- and phospholipids. Boxplot in A) indicates changes in the sum of measured total lipid contents. Lipid composition of C) Monogalactosyldiacylglycerol (MGDG) and D) Digalactosyldiacylglycerol (DGDG) molecular species. Data represent mean values of 5 plants per line. Error bars represent ± SD. Significance of differences between wildtype and mutant lines was analyzed by Student’s t-test: p-value ≤0.05:* (Supplemental File 1).

MGDG and DGDG represent the two most abundant glycerolipids in Arabidopsis leaves (Figure 8A). Interestingly, a closer inspection of the contents of the two MGDG molecular species 34:6 and 36:6 – which are indicative for either plastid-generated MGDG (34:6) or ER-born MGDG (36:6) - revealed clear differences between wild types and overexpressors. Wild-type plants exhibited about 85 mol% of 34:6 MGDG, while leaves from FAX1 Ox#2 and Ox#4 plants accumulated only 75 and 62 mol% of this MGDG species, respectively (Figure 8C). In marked contrast to this, wild type contained only 10 mol% of the MGDG molecular species 36:6, while the FAX1 overexpressor plants Ox#2 and Ox#4 accumulated 21 and 30 mol% of 36:6 MGDG, respectively (Figure 8C). The relative levels of 34:6 and 34:3 DGDG in wild type leaves were higher than in correspondingly grown FAX1 Ox#2 and Ox#4 plants. 34:6 DGDG in wild type amounted at 6.5 mol%, while FAX1 Ox#2 and Ox#4 plants contained only 5.6 or 4.6 mol%, respectively (Figure 8D). Eukaryotic ER-produced 36:6 DGDG in wild types amounted to a relative abundance of 61 mol%, while, similar to the increase of 36:6 MGDG, the two FAX1 overexpressors contained 75 and 74 mol% of 36:6 type DGDG, respectively (Figure 8D). In summary, we can conclude that FAX1 overexpressor lines at cold temperatures accumulate ER-produced phospholipids – namely PC and PE – as well as galactolipids (MGDG, DGDG) with DAG backbones from the eukaryotic pathway, while the proportions of prokaryotic 34:x galactolipids are reduced.

### FAX1 overexpressor lines show symptoms of impaired photosynthesis at early time points of cold exposure

One marked phenotype of FAX1 overexpressor plants after transfer to cold conditions is the appearance of wilted, decayed leaves after some weeks of growth (Figure 6F, H). However, degradation of the FAX1 protein and first changes of the lipid composition are already observed after a few days at 4°C (Figure 1F and Supplemental Figure 1). To search for additional responses, we quantified photosynthetic parameters after short and longer exposure to 4°C. To this end we grew all plants for 28 days at 21°C prior to transfer to at 4°C. At the beginning of the transfer, and after one or six weeks of growth at 4°C, we quantified the photosynthetic performance by measuring the following parameters: the ratio of variable- to maximal fluorescence (*F*_v_*/F*_m_), PSII efficiency Y(II), non-photochemical quenching Y(NPQ), non-regulated quenching Y(NO), and the rate of electron transport (ETR). This comprehensive analyses have been done using the pulse-amplitude-modulation (PAM) fluorometry method (Schreiber, 2004).

Prior to transfer to the cold, all wild type and the two FAX1 overexpressor plants exhibited similar *F*_v_*/F*_m_ ratios (Figure 9A) which ranged around 0.79 (Figure 9B). After onset of cold temperatures, the *F*_v_*/F*_m_ ratio decreased gradually in all three lines. Already after one week, the *F*_v_*/F*_m_ ratio in both FAX1 overexpressor lines was significantly lower than displayed by the wild type, and after six weeks in the cold, wild type plants showed a *F*_v_*/F*_m_ ratio of 0.73, while both mutants exhibited a *F*_v_*/F*_m_ ratio of about 0.68 (Figure 9A, B). This altered *F*_v_*/F*_m_ ratio in the cold is not reflected by an increased NPQ, although FAX1 Ox#2 plants showed a slightly increased NPQ, after 7 days in the cold. However, this was not found for FAX1 Ox#4 plants (Figure 9) and even after 5 weeks in the cold, NPQ in all three lines was similarly decreased (Figure 9C). The photosynthetic quantum yield Y(II) of all three plant lines was similar when grown under control temperature (Figure 9C). In contrast, both FAX overexpressors showed after 7 or 42 days in the cold a decreased Y(II) and an increased energy dissipation via non-regulated quenching Y(NO), which was especially pronounced after 42 days (Figure 9C). While the chloroplastic electron transport rate (ETR) of all three plant lines was similar at the beginning of the cold treatment, the ETR in both FAX1 overexpressor plants was after 7 days at 4°C markedly lower, when compared to corresponding wild type plants (Figure 9C).

**Figure 9:**
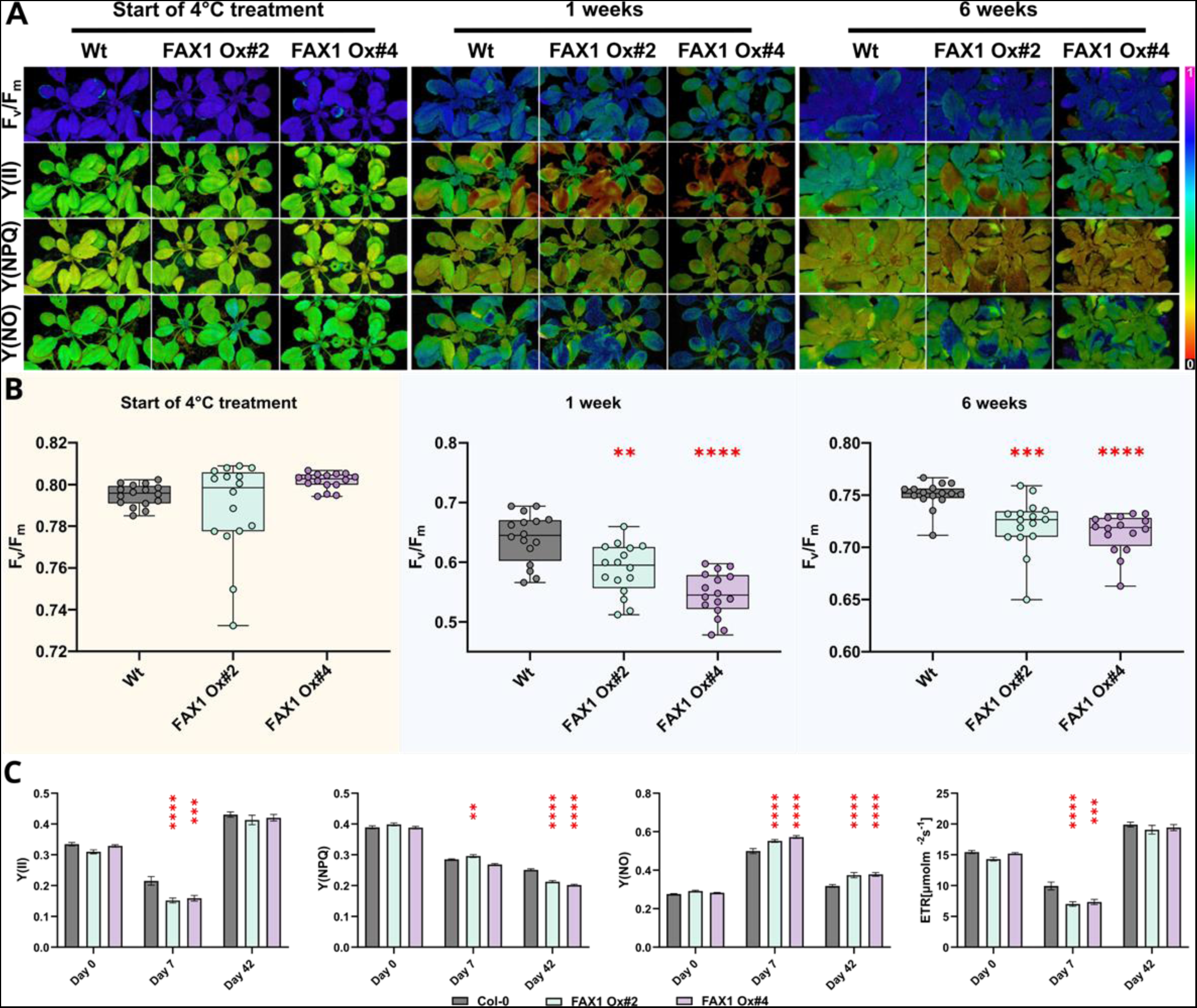
Cold-dependent PSII alterations in FAX1 over expressor lines. After 28 days of growth under short-day (10/14 hours; 110 PAR) conditions at RT, plants (Col-0; FAX1 Oex#2; FAX1 Oex#4) were shifted to a 4°C short-day (110 PAR) chamber. Pulse-Amplitude-Modulation (PAM) induction curve measurements at 110 PAR were performed on day 0 as well as after 7 days and 42 days of the cold treatment. A) Representative PAM images are depicted for PSII capacity (F_v_/F_m_), yield (Y(II)), the quantum yield of light-induced non-photochemical fluorescence quenching (Y(NPQ)), and quantum yield of nonregulated energy dissipation (Y(NO)). B) F_v_/F_m_ determination of the three genotypes at the start of the cold treatment, after one and five weeks. C) Induction curves were generated until a steady-state phase of Y(II), Y(NPQ); Y(NO), and electron transport rate (ETR [µmol m^-2^ s^-1^] were reached. The data shown represents this steady-state after 600 seconds. N = 16; Mean; ± SEM; p-value: (one-way ANOVA): * = 0.033; ** = 0.0021; *** = 0.0002; **** = 0.0001 (Supplemental File 1).

### FAX1 overexpressor and *rbl11* mutants exhibit signs of reactive oxygen species (ROS) accumulation

Several systemic changes in FAX1 overexpressors and *rbl11* mutants occur quite rapidly after transfer to cold conditions. For example, first changes in the lipid composition of both types of mutants are already present after two weeks at 4°C (Figure 5 and Supplemental Figure 1) and alterations in photosynthetic parameters of FAX1 overexpressors are already present after one week of growth at low temperature (Fig. 7). Given that both types of mutants contain higher FAX1 abundance than present in wild types (Figures 1 and 4) and show similar shifts in lipid biosynthesis (Figure 5 and Supplemental Figure 1) we searched for further similarities in their molecular responses after transfer into cold conditions.

The accumulation of both, anthocyanins and reactive oxygen species (ROS) can be taken as early responses upon onset of abiotic stress stimuli (Chalker-Scott, 1999; Baxter et al., 2014). When grown at 21°C, *rbl11* plants and the two FAX1 overexpressor lines contain similar levels of anthocyanins as wild type plants (Figure 10A). In contrast, when exposed to 4°C for only one week, *rbl11* mutants and FAX1 overexpressors accumulated significantly more anthocyanins than the correspondingly grown wild types (Figure 10A). The levels of the transcript coding for the protein BAP1, which indicates the cold-induced occurrence of ROS (Yang et al., 2007; Zhu et al., 2011), was unchanged between the plant lines when grown at 21°C (Figure 10B). Similar to the anthocyanin accumulation, *BAP1* mRNA accumulated to much higher extents in both types of mutants after one week of growth at 4°C, than in wild type plants (Figure 10B).

**Figure 10:**
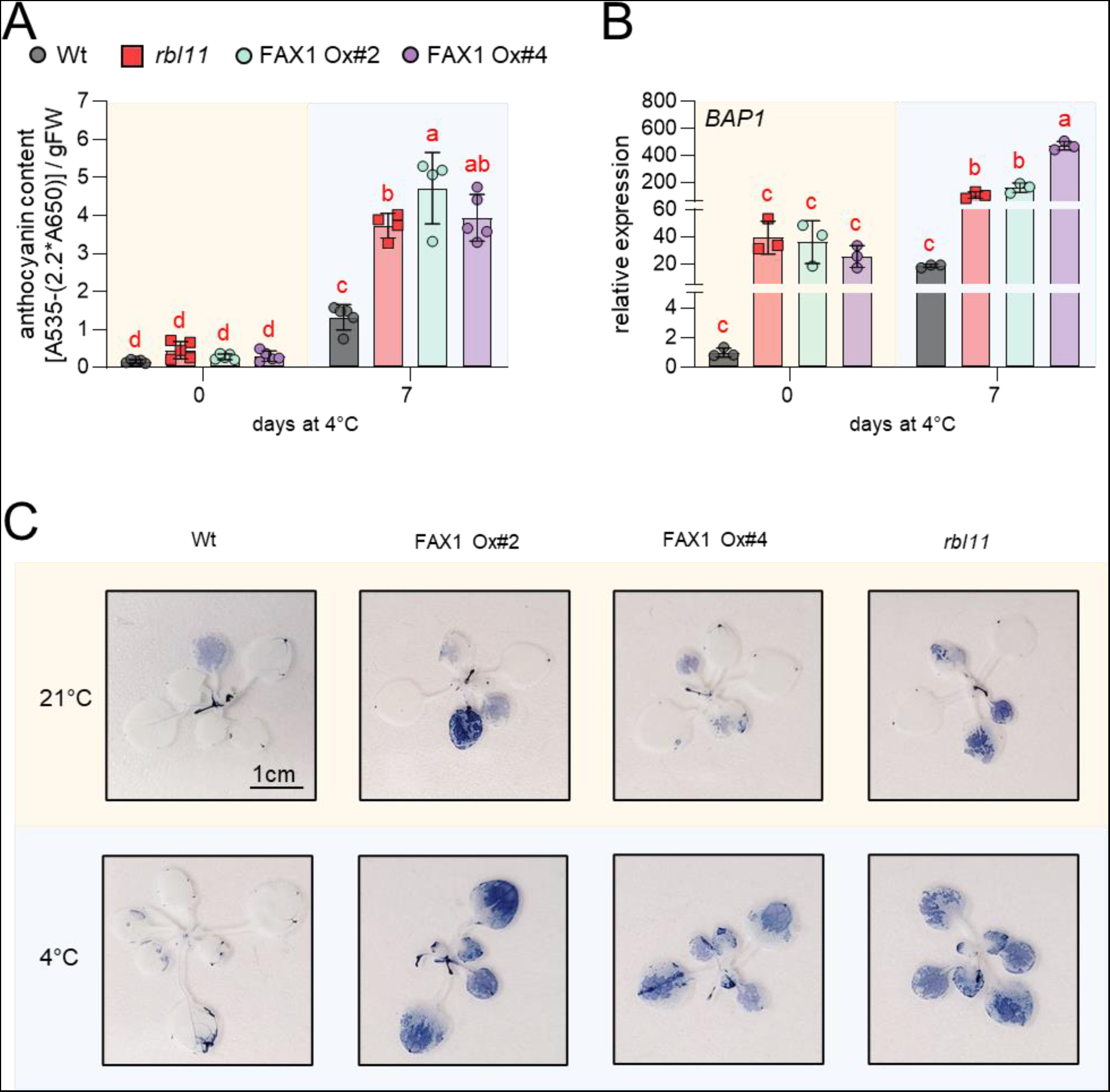
Anthocyanin and reactive oxygen species (ROS) accumulation in *rbl11* loss of function mutants and FAX1 overexpressors at low temperatures. A) Impact of cold on anthocyanin accumulation under ambient conditions and after 7 days at 4°C. B) Relative transcript levels of *BAP1* during standard conditions and after 7 days at 4°C. C) NBT staining of O_2_- accumulation. Plants were grown for 2 weeks under control conditions and subsequently cultivated at 4°C for 4 days. Error bars in A) and B) represent ± SD. Letters displayed over the error bars indicate significant differences analyzed by two-way ANOVA followed by Tukey’s multiple comparisons test (p-value <0.05; (Supplemental File 1)).

To corroborate the differences in cold-induced ROS levels in wild types and the two types of mutants, we analyzed the relative increase in leaf superoxide by nitroblue-tetrazolium (NBT) staining (Doke, 1983; Hoffmann et al., 2013). NBT staining of wild-type tissue at the start of cold treatment and after four days at 4°C does not result in a detectable increase in superoxide (Figure 10C). However, both types of mutants showed a stronger NBT staining after four days at 4°C than at the beginning of cold treatment (Figure 10C).

### FAX1 overexpressor plants exhibit altered levels of metabolites when exposed to cold temperatures

To gain additional insight into metabolic causes for the impaired cold- and frost tolerance pattern of FAX1 overexpressor lines, we analyzed the levels of 71 primary metabolites via GC-MS. Thus, wild-type plants and both FAX1 overexpressor lines were grown for 28 days at 21°C prior to growth at 4°C for two weeks.

When cultivated at 4°C for two weeks, the FAX1 overexpressor lines showed the largest differences in their metabolic readjustment compared to correspondingly grown wild-type plants (Figure 11, Supplemental Table 4). The two FAX1 overexpressor lines contained nearly doubled levels of β-alanine, asparagine, serine and ethanolamine, 1.7-fold increased amounts of glycerol-phosphoglycerol and 1.7-fold more malate when compared to wild types (Figure 11). Similar to malate, three further intermediates of the tricarboxylic acid cycle, namely citrate, fumarate and succinate were also significantly higher in cold treated FAX1 overexpressors than in wild type plants (Figure 11). In addition, cold treated FAX1 overexpressors contained about 38% more proline, about twice as much putrescine and 2.3-fold more spermidine when compared to corresponding wild types, whereas the trehalose concentration in FAX1 Ox#2 plants was substantially lower, compared to wild types (Figure 11).

**Figure 11:**
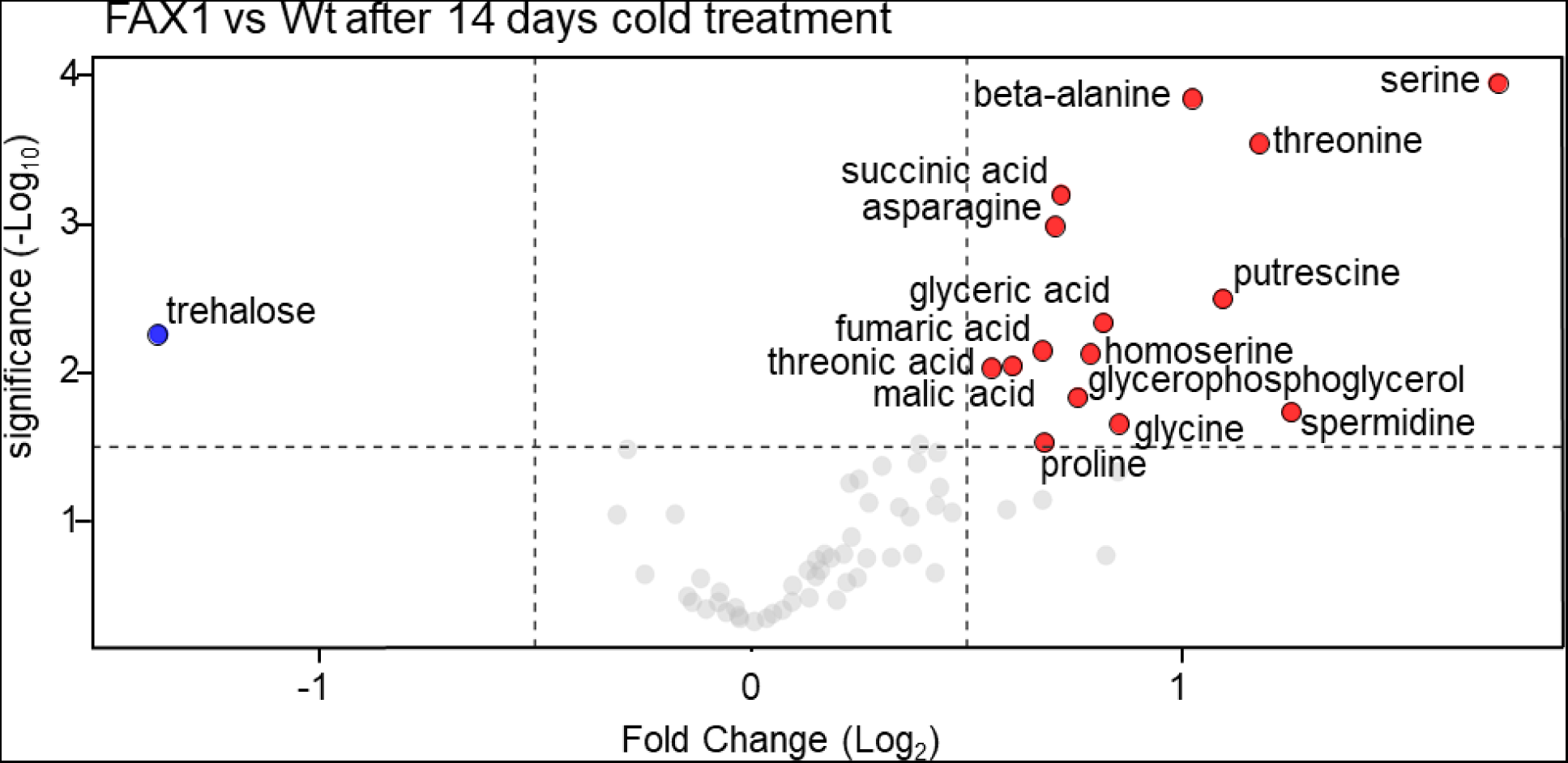
Volcano plot of the metabolic differences measured between Wt and FAX1 Ox#2 rosette leaves, cold treated for 14 days at 4°C. Blue dots represent decreased and red dots increased metabolites with a log_2_ fold change ≥ 0.5 and p-value ≤ 0.05. The complete data set is available in Supplemental Table 4. Significant differences between the 21°C and 4°C treatments were analyzed using a t-test (Supplemental File 1).

## Discussion

The dynamic modification of plant organelle proteomes is mandatory to achieve new homeostatic levels allowing to cope with challenging environmental conditions (Taylor et al., 2009). As seen in many systems, cellular mRNA level and the respective protein amounts do not necessarily correlate to a high degree (Gygi et al., 1999; Greenbaum et al., 2003; Koussounadis et al., 2015). Thus, other factors are also important for controlling protein abundance. The chloroplast envelope proteome undergoes substantial remodeling in response to changes in light or temperature conditions (Knopf et al., 2012; Nishimura et al., 2016; Adam et al., 2019; Mielke et al., 2020) and it can thus be expected that proteases must play a role for this.

Interestingly, cold-induced changes in the proteome are accompanied by modifications of membrane lipids. Latter process maintains membrane fluidity at low temperatures and stabilizes membrane integrity (preventing rigidification) to ensure proper organelle function (Moellering et al., 2010; Zheng et al., 2011; Barnes et al., 2016; Barrero-Sicilia et al., 2017; Guo et al., 2018). For lipid biosynthesis, especially for the generation of phospholipids, fatty acid export from the chloroplast is mandatory. and the FAX1 protein is the best characterized chloroplast envelope located protein presumed to be involved in this transport process so far (Li et al., 2015; Li et al., 2016; Xiao et al., 2021).

In Arabidopsis FAX1 abundance rapidly decreases after transfer to cold temperatures while other envelope proteins increase (Trentmann et al., 2020). Taking the proteins NTT (the chloroplast ATP importer (Tjaden et al., 1998) or MEX1 (the chloroplast maltose exporter) (Niittylä et al., 2004) as examples, we demonstrated that the relative changes of these carriers are key to proper cold tolerance (Trentmann et al., 2020). However, in the case of FAX1, it is unknown whether the decreased protein abundance is a controlled process required to tolerate low environmental temperatures. In addition, no protease has been discovered that could mechanistically explain the decreased abundance of selected envelope-associated proteins in the cold.

Compared to *RBL10* and *FtsH11* mRNAs, the *RBL11* transcript accumulates early after cold exposure (Figure 1A). This correlation is a first indication of a specific molecular interaction between RBL11 and FAX1, and a physical contact between these two proteins, which is a prerequisite for FAX1 degradation, was confirmed by BiFC analysis (Figure 2). The suggestion of a specific effect of RBL11 on FAX1 abundance in the cold is supported by two observations. First, cold-induced FAX1 degradation did not occur in *rbl11* mutants (Figure 1E), whereas *rbl10* or *ftsh11* mutants degraded FAX1 in the cold similarly to wildtype plants (Figure 1E). Second, dexamethasone-induced expression of RBL11 leads to a decrease in FAX1 protein (Figure 4C).

In general, the ability of RBL11 to degrade intrinsic membrane proteins with multiple transmembrane domains, such as FAX1 (Li et al., 2015), is consistent with the properties of other rhomboid proteases (Erez and Bibi, 2009). However, since dexamethasone-induced expression of RBL11 leads to FAX1 degradation only at cold temperature and not at 21°C (Figure 4C), we propose a so far unknown post-translational modification of RBL11 and/or FAX1, which is a prerequisite for catalytic protease activity (Figure 4C). The observation that RBL11 activity might also be involved in the dynamic change TGD5 and TGD2 (Table 1), two components of the TGD complex involved in the unidirectional ER to plastids import of eukaryotic lipids (Xu et al., 2010; Li-Beisson et al., 2017), point to a central function of this protease in modification of plant lipid homeostasis under challenging temperature conditions. Notably, RBL10, the closest RBL11 homologue, has also been shown to affect lipid metabolism as it interacts with the ACYL CARRIER Protein4 and modulates MGDG biosynthesis (Lavell et al., 2019; Xu et al., 2023). Because *rbl11* mutants exhibit a shift toward eukaryotic lipid biosynthesis (which is due to decreased FAX1 abundance but not observed in wild types, Figures 3 and 6, Supplemental Figure 1), we hypothesize that *rbl11* cells attempt to reduce the unintended stimulation of eukaryotic lipid biosynthesis by downregulating the core plastid lipid importer TGD (Table 1). This assumption is supported by the observation that Arabidopsis, as a 16:3 plant, normally stimulates the plastid membrane lipid pathway after exposure to cold conditions (Li et al., 2015; Yu et al., 2023).

Apart from FAX1 and TGD components, which act as substrates for RBL11, it is worth mentioning that the protochlorophyllide oxidoreductase PORA strongly accumulates in *rbl11* mutants (Table 1). PORA is responsible for the stromal conversion of protochlorophyllide to chlorophyllide and it was shown, that increased protochlorophyllide oxidoreductase activity leads to ROS formation (Pattanayak and Tripathy, 2011). Thus, RBL11 might not only contribute to the regulation of FAX1 activity but also influence envelope-located mechanisms preventing cold-induced ROS formation.

*rbl11* mutants exhibit in cold conditions both, higher FAX1 protein levels than observed in wild types (Figure 1E) and a shift of their lipid composition towards accumulation of phospholipids (Figure 5). The latter observation is in accordance with both, the function of FAX1 as a chloroplast to ER fatty acid (FA) export protein (Li et al., 2015; Li et al., 2016; Li-Beisson et al., 2017; Takemura et al., 2019; Tian et al., 2019) and the function of the ER as the cellular site of phospholipid biosynthesis (Li et al., 2015; Li et al., 2016; Li-Beisson et al., 2017; Takemura et al., 2019; Tian et al., 2019). In fact, apart from a stimulation of the eukaryotic pathway of lipid biosynthesis in the cold, *rbl11* plants and FAX1 overexpressors share further similarities, e.g., impaired tolerance to low temperatures and frost (Figures 1, 2 and 7), and increased levels of anthocyanins, ROS and *BAP1* transcripts in the cold when compared to wild types (Figure 10A-C). Latter changes are independent molecular markers for a reinforced stress situation in mutants (Chalker-Scott, 1999; Choudhury et al., 2017)

Although in sum these similarities are indicative for an important function of the down regulation of FAX1 during cold tolerance, a detailed analysis of the impact of FAX1 on this process cannot be made in the *rbl11* mutants, since RBL11 acts on several inner envelope proteins, which triggers pleiotropic effects in *rbl11* plants (Supplemental Tables 1 and 3, and Table 1). Thus, the impact of FAX1 on cold and frost tolerance was analyzed using FAX1 overexpressors. Although the relative abundancies of *FAX1* mRNA and protein differed substantially between Ox#2 and Ox#4 mutants, it turned out that the physiological responses of the two plant lines were very similar (Li et al., 2015 and below). Accordingly, these lines are suitable to search for a potential impact of a cold-induced down regulation of FAX1 levels for tolerance to low temperatures.

The observation that FAX1 overexpressor leaves exhibited higher levels of ER-derived membrane lipids (PE, PC) when exposed to cold temperature (Figure 8 and Supplemental Figure 1B) indicates that the cold-induced decrease of FAX1 in wild types (Trentmann et al., 2020) is a limiting factor for synthesis of lipid backbones via the eukaryotic pathway at low temperature. Our assumption that a relative stimulation of lipid biosynthesis in the ER is causative for increased total levels of galacto- and phospholipids in FAX1 overexpressors gains independent support by observations made on mutants lacking the envelope proteins TGD2 or TGD3. Corresponding loss-of-function mutants show decreased levels of MGDG and DGDG with ER-derived DAG-backbones, while ER-borne phospholipid levels are increased (Awai et al., 2006; Lu et al., 2007). In other words, a higher substrate availability such as acyl residues at the ER might lead to increased levels of membrane lipids with ER-derived DAG backbones (Figure 12).

**Figure 12:**
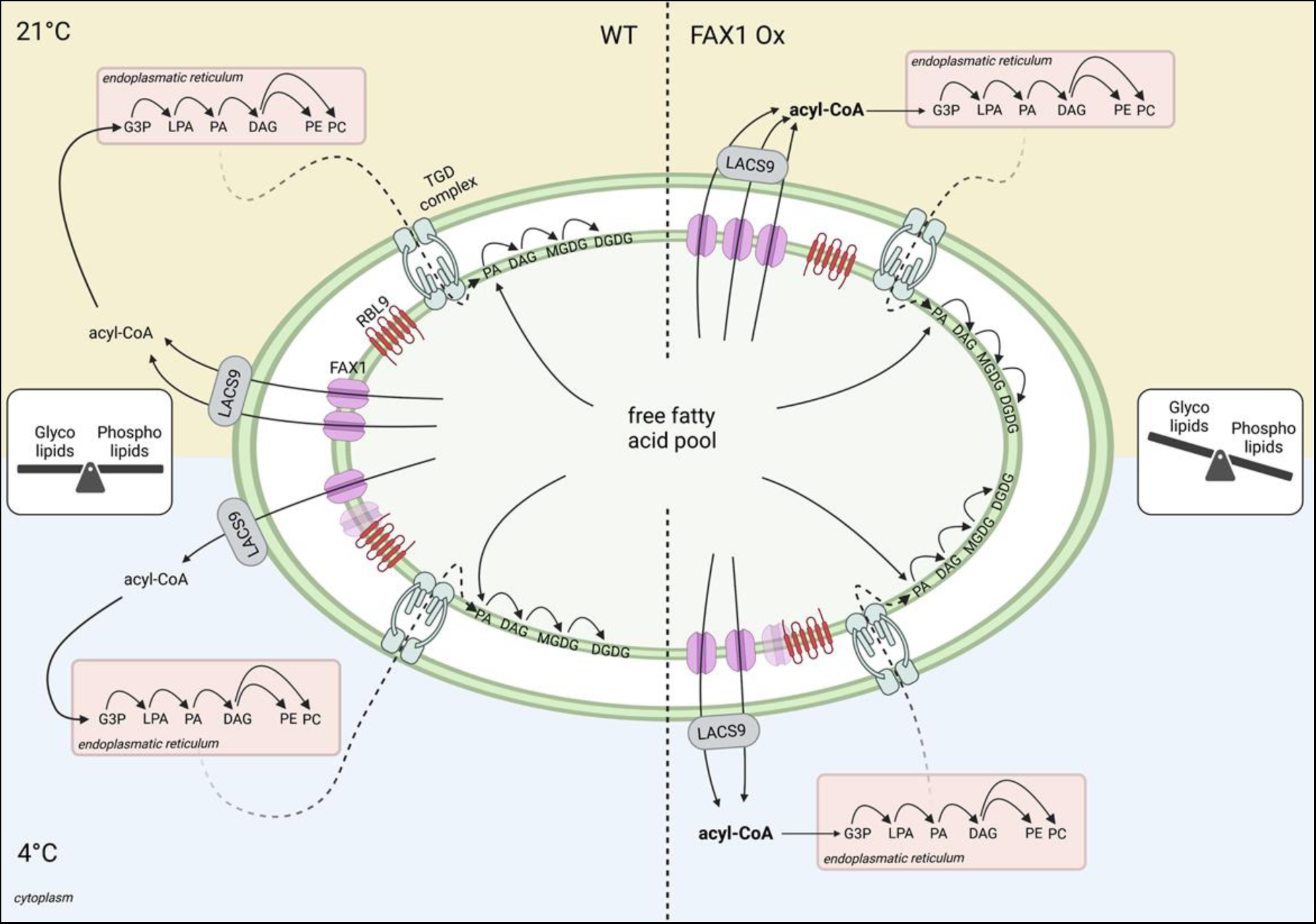
Proposed model for the influence of the fatty acid export protein1 (FAX1) abundance, controlled by the envelope located rhomboid-like protease11 (RBL11), on cold adaptation via balancing the glyco- and phospholipid contents in Arabidopsis. In wild types, the the glyco- and phospholipid levels are balanced to gain ideal conditions for growth and development under diverse environmental conditions. During low temperature, RBL11 interacts with FAX1, which leads to a decrease of FAX1 abundance. The associated reduced export of fatty acids, and concomitted slowdown of the eukaryotic pathway for lipid biosynthesis, seems to be an efficient mechanism for cold acclimation. In FAX1 overexpressing (Ox) plants however, the permanent increased export of fatty acids from the chloroplast, which stimulates lipid biosynthesis in the Endoplasmic Reticulum, leads to a shift to phospholipid synthesis. In FAX1 Ox lines, RBL11 is unable to decrease FAX1 protein abundance, and the disturbed glyco- to phospholipid ratio impairs the efficient acclimation to cold temperatures, results in cold- and frost sensitive mutants.

MGDG represents the most abundant chloroplast-located galactolipid (Dorne et al., 1990; Kobayashi, 2016) and in the 16:3 plant Arabidopsis, two main molecular species occur i.e., plastid derived 34:6 type MGDG and, as ER derived, 36:6 type MGDG (Boudière et al., 2014). When compared to wild types, Ox#4 mutants grown at 21°C exhibit a relative decrease of 34:6 type MGDG, which is nearly balanced by a corresponding increased level of 36:6 type MGDG (Figure 8C), and the proportion of 34:6 DGDG is decreased, while 36:6 DGDG is increased when comparing wild type with Ox#4 plants grown at 4°C (Figure 8D). Thus, the latter observation indicates that a stimulated fatty acid export in Ox#2 and Ox#4 plants leads to a shift from plastid-derived to ER-derived galactolipids. Such flexible shift of lipid biosynthesis from plastids to the ER has already been observed for mutants lacking the plastid glycerol-3-phosphate acyltransferase (ACT) activity (Falcone et al., 2004; Lusk et al., 2022), and also in *act1* mutants (synonymous: *ats1*) a stimulation of membrane lipid biosynthesis in the ER largely compensates for the impaired prokaryotic lipid biosynthesis (Kunst et al., 1988). The observation that FAX1 overexpressors not only exhibit an increased ratio of 36:6 to 34:6 type MGDG (Figure 8C), but also increased ratios of 36:6 to 34:6-, and 36:6 to 34:3 type DGDGs (Figure 8D) further underlines the shift from prokaryotic to eukaryotic lipid biosynthesis in these lines. Finally, the increase of the relative proportions of the two most abundant phospholipids, PC and PE, supports the conclusion that the eukaryotic lipid biosynthesis pathway is stimulated in FAX1 overexpressors.

The analyses of the molecular responses of the FAX1 mutants to cold and freezing conditions showed that these changes are associated with a strongly impaired ability to resist low environmental temperatures, as evidenced by an increased number of wilted leaves in the cold and a decreased ability to recover from freezing (Figures 2 and 7). A similar gradual decay of leaves was also seen for other Arabidopsis mutants exhibiting decreased amounts of polyunsaturated fatty acids in membrane lipids (Miquel et al., 1993) and of course for the *rbl11* mutant (Figure 1). It seems likely, that the increased number of wilted leaves of FAX1 overexpressors when grown at 4°C (Figure 6E and F) is the result of an impaired photosynthetic performance (Figure 9A,C).

The exact reasons for these marked effects are not clear, but it was shown that changes in the composition of membrane lipids, induced by modifications of different lipid biosynthesis genes, affect photosynthetic properties (Botté et al., 2011; Kobayashi, 2016; Gao et al., 2020). Thus, it seems conceivable that the altered chloroplast membrane composition observed in FAX1 overexpressors (Supplemental Figure 1 and Figure 8) is causative for the impaired photosynthetic performance. Indeed, the decreased F_v_/F_m_ ratio of FAX1 overexpressor plants (Figure 9A,B) is an indicator of impaired PSII function (Murchie and Lawson, 2013) and a temperature stressed PSII can lead to ROS production (Pospíšil, 2016), as observed in FAX1 overexpressor plants (Figure 10C-F). However, it cannot be excluded that the marked effects of RBL11 mutation or FAX1 overexpression on PC and PA levels (Figures 5C,D and 8A,B) also contribute to ROS accumulation (Figure 10). This is because the signaling molecule PA has been shown to bind directly to the plasma membrane NADPH oxidoreductase RBOHD, leading to the activation of this enzyme. Accordingly, this process stimulates O_2_^-^ production, which ultimately leads to H_2_O_2_ accumulation (Zhang et al., 2009).

Wild-type plants and FAX1 overexpressor lines exhibit a quite similar metabolite pattern when grown at 21°C. In contrast, the metabolite composition of FAX1 Ox plants during growth at 4°C is to some degree indicative for reinforced cold stress. For example, the comparably high accumulation of polyamines in form of putrescine and spermidine, of amino acids like proline and asparagine, or the accumulation of the non-proteinogenic amino acid β-alanine (Figure 11) represent independent evidences for a pronounced metabolic response to cold temperatures (Alet et al., 2011; Liang et al., 2013; Zhang et al., 2016; Marco et al., 2019; Parthasarathy et al., 2019). β-alanine is not only a metabolic indicator for abiotic stress (Parthasarathy et al., 2019), it also acts as precursor for CoA synthesis and is thus required for fatty acid and phospholipid biosynthesis (Perrett et al., 2017). Therefore, β-alanine accumulation is coincident with higher total levels of galacto- and phospholipids in FAX1 overexpressors in the cold (Figure 11A). Under many stress conditions, asparagine and proline levels change similarly (Curtis et al., 2018). While the exact function of asparagine during stress is unclear, the role of proline as a general stress metabolite is well established (Liang et al., 2013; Ghosh et al., 2022). For example, proline is able to diminish PS_II_ defects caused by rising ROS levels (Alia and Mohanty, 1997) and because cold treated FAX1 overexpressors exhibit defects in PS_II_ activity (Figure 9A-C), the induce proline accumulation might represent a process to tune down such negative effects.

In summary, we propose a model in which the envelope protease RBL11 is responsible for the degradation and down-regulation of FAX1 in cold-treated Arabidopsis plants (Figure 12). This process represents a previously hidden molecular response that is critical for optimal low temperature acclimation. Most likely, the initial cold response in wild-type plants is represented by a preference for the prokaryotic lipid synthesis pathway to protect chloroplast membranes and the photosynthetic machinery from cold damage. Since the corresponding *FAX1* mRNA is not decreased in cold-treated wild-type plants (Figure 6A), the decreased FAX1 protein level is most likely due to a specific effect of RBL11. In near future it will be interesting to search for factors that activate RBL11 and/or convert selected membrane proteins on the inner envelope proteins into specific substrates (Figure 12). It seems worth to mention, that the cold-induced degradation of FAX1 further supports findings on the importance of various chloroplast envelope associated processes for plant cold and frost tolerance (Moellering et al., 2010; Barnes et al., 2016; Guan et al., 2019; Schwenkert et al., 2023).

## Material and Methods

### Plant cultivation and growth conditions

Arabidopsis (*Arabidopsis thaliana*) ecotype Columbia (Col-*0*) and transgenic plants were sown on standard soil (type ED73, Einheitserde Patzer; Sinntal-Altengronau, Germany) with 10% (v/v) sand, stratified at 4°C for 48 h and then grown under short-day regime (10 h light/14 h dark) at 60% relative humidity and 120 µmol m^−2^ s^−1^ light intensity at 21°C, representing standard growth conditions. For cold treatment, plants were were grown for 21 days at 21°C first and subsequently transferred to a cultivation chamber (Fitotron SGR223, Weis-Gallenkamp Technik, Heidelberg, Germany) and incubated for several days at 4°C while all other parameters were kept constant.

We descripted FAX1 Ox#2 and FAX1 Ox#4 lines earlier (Li et al., 2015). *rbl10* mutants (Lavell et al., 2019) were provided by Dr. Christoph Benning (Michigan State University, Wisconsin, USA), *ftsh11* and *rbl11* mutants (Knopf et al., 2012; Adam et al., 2019) were provided by Dr. Zach Adam (Hebrew University, Jerusalem, Israel).

For RNA extraction, metabolite and anthocyanin analysis, Arabidopsis rosette leaves were collected five hours after onset of light, transferred immediately into liquid nitrogen and stored at −80°C until preparation. Leaf material for chloroplast envelope isolation was used directly after harvesting one hour before the start of illumination. For lipid isolation, rosette leaves were collected and directly transferred in a glass tube containing boiling water.

### Generation of RBL11 overexpressor lines

The cloning steps to generate the dexamethasone-inducible RBL11 overexpression lines were performed using S7 Fusion Polymerase™ (MD-S7-100, Mobidiag, Espoo, Finland). The sequence of BirA-HA was first amplified from the Vector pcDNA3.1 MCS-BirA(R118G)-HA (Roux et al., 2012) using gene-specific primers and ligated C-terminally of *RBL11* in the pBSK vector (Short et al., 1988) by overlap extension PCR. The *RBL11*-BirA-HA construct was amplified using gene-specific primers with *attB*-sites attached and inserted first into pDONR and then into the destination vector pTA7001-DEST (Aoyama and Chua, 1997) via the Gateway Cloning system.

Wildtype Arabidopsis plants were transformed via floral dip using *Agrobacterium tumefaciens* (GV3101). Positive transformed plants were selected by hygromycin selection (Harrison et al., 2006) and confirmed by Western Blot analysis using an HA-antibody after induction of leaves with 30µM dexamethasone. Sequences of gene-specific primers, which were used for cloning, are provided in Supplemental File 2

### Protein extraction from RBL11 overexpressor plants

Prior to protein extraction, approximately 50 leaves of Arabidopsis wildtype, or RBL11-HA overexpressor plants were cut and placed in 30µM DEX or water, as control. Leaves were incubated on a shaker for 48h in a plant chamber at either 21°C, or 4°C. Preparation of soluble and insoluble (membrane) protein fraction and detection of RBL11-HA using HA antibody via Western Blot was performed as described earlier (Khan et al., 2018).

### Arabidopsis chloroplast envelope preparation

Arabidopsis chloroplast envelope membranes were isolated according to an established protocol (Bouchnak et al., 2018) with few modifications. In a cold room (4°C), before onset of light, 100 to 200 g of rosette leaves were harvested from six-weeks old plants and ground in a Waring blender (three cycles, each of 2 seconds, average intensity) in the presence of grinding medium (Tricine-KOH (20 mM, pH 8.4), sorbitol (0.4 M), EDTA (10 mM, pH 8), and NaHCO_3_ (10 mM), BSA (0.1% (w/v)). The homogenate was filtered through one layer of Miracloth and centrifuged for 2 min at 2,070x*g* at 4°C. The supernatant was discarded, and the sediment was gently resuspended on ice with a soft natural bristle paint brush in washing medium (1x) (Tricine-KOH (10 mM, pH 7.6), sorbitol (0.4 M), MgCl_2_ (2.5 mM), and EDTA (1.25 mM)) with a final volume of the combined chloroplast solutions = 24 ml). 6 ml of the suspension was equally distributed and loaded on top of four continuous Percoll (Sigma Aldrich, Heidelberg, Germany) gradients (containing 50% Percoll / 0.4 M sorbitol, prepared by centrifugation at 38,700x*g*, for 55 min at 4 °C). Loaded gradients were centrifuged for 10 min at 13,300x*g*, 4°C using a swinging-bucket rotor. The intact chloroplasts present in the lower phase were retrieved with a 10 ml pipet. The intact chloroplast suspension was washed twice with 30 ml washing buffer (1x) and centrifuged for 2 min at 2,070x*g* at 4°C. After washing, the purified chloroplasts were lysed by resuspending the sediment in hypotonic medium (MOPS (10 mM, pH 7.8), MgCl_2_ (4 mM), PMSF (1 mM, dissolved in isopropanol), benzamidine hydrochloride hydrate (1 mM), and ε-amino caproic acid (0.5 mM). 3 ml of lysed chloroplasts were loaded on top of two prepared sucrose gradients (4 ml of 0.93 M, 3 ml of 0.6 M and 2.5 ml of 0.3 M sucrose). Gradients were ultracentrifuged for 1 h at 70,000x*g*, 4°C in a swinging-bucket rotor. The yellow band of both gradients (containing the envelope fraction) was retrieved and pooled in one tube. The envelope suspension was washed in 12 ml membrane washing buffer medium (MOPS (10 mM, pH 7.8), PMSF (1 mM), benzamidine hydrochloride hydrate (1 mM), ε-amino caproic acid (0.5 mM) and ultracentrifuged again for 1 h at 110,000x*g*, 4 °C. Supernatants were aspirated by using a water pump. Approximately 100 µl of membrane washing buffer was used to resuspend the envelope sediment. Isolated envelopes were stored in liquid nitrogen until use.

### Bimolecular Fluorescence Complementation for interaction studies

For the cloning of the BiFC constructs, the full-length sequences of *RBL11, FAX1* and *FTSH11* were used. The coding sequences were amplified by PCR using S7 Fusion Polymerase (MD-S7-100, Mobidiag, Espoo, Finland) and inserted first into pDONR and then into the pUBC-cYFP and the pUBC-nYFP vectors via the Gateway cloning system (Grefen et al., 2010). Half of a yellow fluorescent protein (nYFP or cYFP) is thus fused to the C-terminus of *RBL11*, *FAX1*, or *FTSH11*. The resulting constructs were then transformed into *Agrobacterium tumefaciens* strain GV3101. Transient expression in tobacco (*N. benthamiana*) leaves of RBL11, FAX1, and FTSH11, each fused to an nYFP or cYFP, was performed as described (Walter et al., 2004). *Nicotiana benthamiana* leaves were infiltrated through the lower epidermis. After 5 days, leaves were analyzed using a Leica TCS SP5II fluorescence microscope (Leica Instruments, Wetzlar, Germany) (514 nm excitation and 525-582 nm detection of emission through an HCX PL APO 63 × 1.2 W water immersion objective).

Sequences of gene-specific primers, which were used for cloning, are provided in Supplemental File 2.

### Frost recovery experiment

For detection of the ability to recover from frost, a freezing tolerance test was performed according to an established approach (Trentmann et al., 2020; Cvetkovic et al., 2021). Survival rates and the numbers of wilted leaves were documented after 7 days of recovery under standard growth conditions.

### Pulse Amplitude Modulation (PAM) Fluorescence Measurements

Photosystem II parameters at constant light intensities were monitored using an imaging PAM M-Series IMAG-K7 and the ImagingWinGigE V2.56p (WALZ, Würzburg, Germany) software. Induction curve settings were on default with 110 PAR as light intensity, 40 s delay- and 20 s clock-time. Dark adaptation of plants lasted 10 minutes, followed by a 615 s long measurement monitoring PSII capacity (*F_v_/F_m_*), PSII effective photochemical quantum yield (Y(II)), the quantum yield of light-induced non-photochemical fluorescence quenching (Y(NPQ)) and quantum yield of nonregulated energy dissipation (Y(NO)) (Genty et al., 1989; Kramer et al., 2004). All plants analyzed (Col-*0*, FAX1 Ox#2 and FAX1 Ox#4) were grown for 28 days under short-day conditions (10/14 h; 110 µE) at RT before being shifted for the duration of 6 weeks to 4°C (10/14 h, 110 µE). PAM measurements were carried out on the day of the shift to 4°C, after one or six weeks in the cold.

### Determination of anthocyanin content

For anthocyanin quantification 1 ml of extraction buffer composed of H_2_O, propanol and HCl (81:18:1) was added to 100 mg of fine grounded rosette plant material and incubated for 3 min at 95°C, while shaking at 650 rpm and stored over night at RT in full darkness. After centrifugation for 15 min at 12.500 rpm at RT, the supernatant was used for photometric quantification at E1=535 nm and at E2=650 nm. The extinction was determined and corrected (E_corr_=[E_535_-(2.2*E_650_)] / mg FW).

### Metabolomics

Metabolite profiling was performed according to established protocols (Roessner et al., 2001; Lisec et al., 2006; Erban et al., 2007). In brief, from four plants per genotype and growth condition, 50 mg fresh weight (Fw) of ground rosette material was mixed, in a 1.5 ml reaction tube, with 180 µl of cold (−20°C) methanol containing internal standards (10 μl ribitol, 0.2 mg ml^-1^ in water and 10 μl ^13^C-sorbitol, 0.2 mg ml^-1^ in water). After 15 min of incubation at 70°C, the extract was cooled down to room temperature and carefully mixed with 100 μl of chloroform and 200 µl of water. To force phase separation, a 15 min centrifugation step at full speed was performed. Fifty μl of the upper (polar) phase was dried *in vacuo* and stored at −80°C. For derivatization, the pellet was resuspended in 10 μl of methoxyamin-hydrochloride (20 mg ml^-1^ in pyridine) and incubated for 90 min at 40°C. After addition of 20 µl of BSTFA (*N*,*O*-bis[trimethylsilyl]trifluoroacetamide) containing 2.5 μl retention time standard mixture of linear alkanes (n-decane, n-dodecane, n-pentadecane, n-nonadecane, n-docosane, n-octacosane, n-dotriacontane), the preparation was incubated at 40°C for further 45 min.

One μl of each sample was injected into a GC–TOF–MS system (Pegasus HT, Leco, St Joseph, USA). Samples were automatically processed by an autosampler system (Combi PAL, CTC Analytics AG, Zwingen, Switzerland). Helium acted as carrier gas at a constant flow rate of 1 ml min^-1^. Gas chromatography was performed on an Agilent GC (7890A, Agilent, Santa Clara, CA, USA) using a 30 m VF-5ms column with 10 m EZ-Guard column. The temperature of the split/splitless injector was set to 250°C, as well as the transfer line and the ion source. The initial oven temperature (70°C) was linearly increased to a final temperature of 350 °C by a rate of 9°C/min. Metabolites were ionized and fractionated by an ion pulse of 70 eV. Mass spectra were recorded at 20 scans s^-1^ with an *m/z* 50– 600 scanning range. Chromatograms and mass spectra were evaluated using ChromaTOF 4.72 and TagFinder 4.1 software (Luedemann et al., 2008).

### RNA Extraction, cDNA synthesis and qRT-PCR

RNA was extracted from 50 mg of frozen and fine ground rosette leaf material from four biological replicates per genotype and growth condition using the NucleoSpin RNA Plant Kit (Macherey-Nagel, Düren, Germany), according to the manufacturer’s protocol. The synthesis of cDNA from RNA was performed with the qScript cDNA Synthesis Kit (Quantabio, Beverly, MA, USA). Primers used for gene expression analysis via qRT-PCR are listed in Supplemental File 2. *AtUBQ* was used as reference gene for normalization.

### Nitroblue tetrazolium (NBT) staining

For ROS staining, O_2_^-^ was detected by nitroblue tetrazolium staining (Fryer et al., 2002) in whole rosettes of two weeks old Arabidopsis plants cultivated as described above. For cold treatments, plants were transferred to 4°C for four days, while control plants were kept under standard conditions.

### Measurement of galacto-, phospho- and sulfolipids

For the analysis of lipids, plants were cultivated as described above and transferred from standard conditions to 4°C for 2 weeks, 10 weeks, or used directly. From 5 plants per genotype 100 mg fresh weight of rosette leaf material was harvested and immediately placed into a glass tube containing boiling water to prevent degradation of phospholipids through phospholipase D activity. The lipid extraction was performed with chloroform/methanol after deactivation of lipase activities by boiling the tissue in water as described earlier (Gasulla et al., 2013). Lipids were measured by tandem mass spectrometry (Q-TOF 6530 Agilent Technologies) and quantified by MS/MS experiments with internal standards following the strategy developed earlier (Gasulla et al., 2013; Welti et al., 2002).

### Immunoblotting

Per lane, 8 µg of isolated chloroplast envelope protein or 30 µg of freshly prepared protein extract from Arabidopsis leaf material were separated via SDS-PAGE (12%). The proteins in the gel were transferred onto a nitrocellulose membrane by a semi-dry blotting system (TransBlot® Turbo™ Transfer System, BIO RAD, Göttingen, Germany). The membrane was blocked in phosphate-buffered saline plus 0.1% (v/v) Tween 20 (PBS-T) with 3% milk powder for 1 h at room temperature, and then washed three times in PBS-T for 10 min. The membrane was incubated with a polyclonal rabbit antibody raised against FAX1 (Li et al., 2015) over night at 4°C at 1:1000 dilution. After three times of washing with PBS-T for 10 min, the membrane was incubated with a horseradish peroxidase (HPR) conjugated anti-rabbit antibody (Promega, Walldorf, Germany) diluted 1:10.000 in PBS-T with 3% milk powder for 1 h. The immunoreaction was visualized by chemiluminescence using ECL Prime Western blotting reagent (GE Healthcare, Karlsruhe Germany) and a Fusion Solo S6 (Vilber-Lourmat, Eberhardzell, Germany).

### Peptide Mass spectrometry

Enriched envelope fractions were precipitated in 80% acetone, digested in solution using Lys-C and trypsin, and resulting peptides were desalted as previously described (Hammel et al., 2018). Peptide mass spectrometry was performed using a nanoUHPLC-IM-MS system (nanoElute coupled to timsTOF Pro2, Bruker Daltonics, Bremen, Germany). Samples were directly loaded on a 25 cm, 75 µm ID, 1.6 µm particle size, C18 column with integrated emitter (Odyssey/Aurora ionopticks, Melbourne, Australia) set to 50 °C, and peptides were separated under a flow rate of 0.3 µl/min using buffers A (water, 0.1% formic acid) and B (acetonitrile, 0.1% formic acid). The gradient employed ramped from 2% B to 25% B within 67 min, then to 37% B within 10 min, followed by washing and equilibration steps. The MS was operated in positive mode, electrospray voltage was set to 1.4 kV and spectra were recorded from 100 – 1700 m/z. A total of 10 MS/MS PASEF ramps (1/K_0_ 0.6 – 1.43 V*s/cm^2^) with 100 ms duration were acquired per cycle, and target intensity for MS/MS was set to 14500 whereafter the precursors were excluded from fragmentation for 0.4 min.

### Protein Identification and Quantification

Acquired data were searched against the Uniprot protein sequences for *Arabidopsis thaliana* (UP000006548) using the FragPipe v19.1 processing pipeline choosing the default LFQ-MBR workflow with minor modifications: peptide length was set to a minimum of 6 amino acids, missed cleavages to 3, normalization of intensities across runs was omitted. Mass spectrometry raw and processed data have been deposited at the ProteomeXChange Consoritum via the PRIDE partner repository (Perez-Riverol et al., 2021) with the dataset identifier PXD041219.

### Data normalization and missing value imputation

Prior to computation of protein-level statistics, replicate groups were normalized using the median-of-ratios method (Anders and Huber, 2010). Subsequently, we computed global variance estimates and local gene-wise mean estimates to impute missing data points as independent draws from normal distributions. When a replicate group did not contain any measurement, the normal distribution was centered at an intensity corresponding to the 5% quantile of all intensities. Proteins were excluded from statistical analysis according to two filter criteria. First, proteins were considered ineligible for downstream analysis if there was no biological replicate group with at least one reading. Second, only proteins reported to be localized to the envelope membrane were considered, either by association with matching Gene Ontology terms retrieved from UniProt or by entry in the manually curated AT_CHLORO database (Ashburner et al., 2000; Bruley et al., 2012; Consortium, 2021).

### Statistical Analyses

Statistical analyses regarding the proteomic data were based on log2 transformed imputed values. According to the experimental design which consisted of two factors (genotype and treatment) at each of two levels (wild type/mutant and normal/cold), changes in protein abundance were evaluated using two-way analysis of variance, ANOVA (Fisher, 1925). To account for multiple hypothesis testing,we controlled the False Discovery Rate (FDR) by computing q-values based on the ANOVA p-values as previously described by Storey (Storey and Tibshirani, 2003; Storey et al., 2004). All calculations were carried out using the FSharp.Stats library for statistical computing (Venn et al., 2023). Changes were considered to be significant if a q-value of 0.05 was not exceeded.

For statistical analysis of the numerical data GraphPad Prism 9 and Microsoft Office Excel were used. Significant differences between two groups were analyzed by two-tailed Student’s *t-*test. The software Shiny application (https://houssein-assaad.shinyapps.io/TwoWayANOVA/) was used for letter-based representation of all pairwise comparisons using popular statistical tests in two-way ANOVA. Statistical data is provided in Supplemental File 1.

### Accession numbers

Sequence data from this article can be found in the ARAMEMNON GenBank data library (http://aramemnon.uni-koeln.de/). *FAX1* (*At3g57280*), *FTSH11* (*At5g53170*), *RBL10* (*At1g25290*), *RBL11* (*At5g25752*), *BAP1* (*At3g61190*).

## Supporting information

John_et_al_

## Acknowledgments

Work in the labs from ML, H-HK, AF MS, TM and HEN was supported by the Deutsche Forschungsgemeinschaft (DFG) within the SFB/Transregio (TRR) 175, The Green Hub.

## Notes

https://www.proteomexchange.org

### Competing Interest Statement

The authors have declared no competing interest.

### Summary of Updates

Figure 2 and Figure 3 have been revised.

