## Supplementary material for "Degradation of Fatty Acid Export Protein1 by Rhomboid-Like Protease11 Contributes to Cold Tolerance in Arabidopsis": John_et_al_

Supplemental File 1: Primers

| BiFC constructs; Figure 2 | |
|---|---|
| **Primer** | **Sequence** |
| *At*RBL11_fwd_attB | GGGGACAAGTTTGTACAAAAAAGCAGGCTTAATGTCTCAGCTTCTTCACCT |
| *At*RBL11_rev_attB | GGGGACCACTTTGTACAAGAAAGCTGGGTACGAGGACTTTTTAACGTCTTG |
| *At*FAX1_fwd_attB | GGGGACAAGTTTGTACAAAAAAGCAGGCTTAATGGCTTCACAAATCTCTCA |
| *At*FAX1_rev_attB | GGGGACCACTTTGTACAAGAAAGCTGGGTAGTATGAAGGACTAGTCGCAG |
| *At*FtsH11_fwd_attB | GGGGACAAGTTTGTACAAAAAAGCAGGCTTAATGTCTTCTTCTACTCTTCAAG |
| *At*FtsH11_rev_attB | GGGGACCACTTTGTACAAGAAAGCTGGGTATTAGGCCAATACCAAGTCTC |
| qRT PCR; Figure 1A; Figure 4A,B,C | |
| *At*BAP1_RT_fwd | ATCGGATCCCACCAGAGATTACGG |
| *At*BAP1_RT_rev | AATCTCGGCCTCCACAAACCAG |
| *At*UBQ10_RT_fwd | GGCCTTGTATAATCCCTGA TGAATAAG |
| *At*UBQ10_RT_rev | AAAGAGATAACAGGAACGGAA ACATAG |
| *At*FAX1_RT_fwd | CCAAAGCCGATCGTTTCCAT |
| *At*FAX1_RT_rev | AAGTCGTGGATCTTTGCTGC |
| *At*FTSH11_RT_fwd | TGTGGGATGAGTGAGGCAAT |
| *At*FTSH11_RT_rev | GTGTAGCTGCTTCTCATGCC |
| *At*RBL10_RT_fwd | CTGAGAGTTTAGGTGGCCCA |
| *At*RBL10_RT_rev | TCATTGCCACCTCTGACCAT |
| *At*RBL11_RT_fwd | GCTTCTTCACCTTCACCGTC |
| *At*RBL11_RT_rev | GCTCGCTTTTGTGGTTTTCG |
| *RBL11*-HA overexpression lines | |
| BirA-HA_fwd | AGTCCTCGAGGACAACACCGTGC |
| BirA-HA_rev | TGCATGCCTGCGTAATCCGGTACA |
| RBL11-BirA-HA_fwd_attB | GGGGACAAGTTTGTACAAAAAAGCAGGCTTAATGTCTCAGCTTCTTCAC |
| RBL11-BirA-HA_rev_attB | GGGGACCACTTTGTACAAGAAAGCTGGGTACTATGCGTAATCCGGTAC |
